# Lytic IFNγ is stored in granzyme B-containing cytotoxic granules and co-secreted by effector CD8⁺ T cells

**DOI:** 10.1101/2025.01.29.635520

**Authors:** Xuemei Li, Claudia Schirra, Marie-Louise Wirkner, Szu-Min Tu, Chin-Hsin Lin, Meltem Hohmann, Nadia Alawar, Abed Chouaib, Ute Becherer, Varsha Pattu, Jens Rettig, Elmar Krause, Hsin-Fang Chang

## Abstract

Cytotoxic CD8⁺ T cells form immunological synapses with target cells and release effector molecules, including IFNγ, to mediate antitumor immunity. However, the mechanisms by which IFNγ contributes to cytotoxicity remain incompletely understood. Here, we identify a subset of IFNγ stored within GzmB⁺ cytotoxic granules (CGs) in activated mouse and human CD8⁺ T cells, termed ’lytic IFNγ’. Lytic IFNγ is polarized to the synapse and co-secreted with GzmB in both soluble and supramolecular attack particle (SMAP)-associated forms. Mouse CD8⁺ T cells lacking the vesicle priming factor Munc13-4 exhibit impaired both CG and early IFNγ release at the immunological synapse, while prolonged synaptic engagement restores IFNγ secretion. Super-resolution imaging demonstrates that sustained synaptic interactions drive IFNγ secretion at distal membrane sites, suggesting the existence of distinct IFNγ populations with potentially diverse functions beyond lytic IFNγ. These findings uncover an unrecognized mechanism of IFNγ storage and release, underscoring its pivotal role in CD8⁺ T cell-mediated tumor elimination.

## Introduction

Interferon-gamma (IFNγ) is a pivotal cytokine in adaptive immunity, essential for immune regulation and antitumor defense^1, 2, 3, 4, 5, 6, 7^. It enhances the cytotoxic potential of immune cells, including CD8⁺ T cells, natural killer (NK) cells, and other effector cells^4, 8, 9, 10^. While its immunoregulatory functions are well documented, the mechanisms by which IFNγ directly contributes to cytotoxicity remain unclear.

IFNγ promotes antitumor activity through both direct cytotoxic effects and immune modulation. Its expression correlates with T cell cytotoxicity^11, 12^ and is critical for CD4⁺ CAR T cell-mediated tumor clearance^13^. Autocrine and paracrine IFNγ signaling are vital for CD8⁺ T cell function, and disruption of these pathways significantly impairs cytotoxicity^11^. However, IFNγ also exhibits context-dependent immunosuppressive effects, including immune suppression in certain tissues, modulation of T cell exhaustion, and regulation of regulatory T cell differentiation^14, 15, 16^.

CD8⁺ T cells eliminate infected or malignant cells by releasing lytic molecules at immunological synapses^17, 18^. This process involves the polarization and secretion of lytic granules containing perforin, granzyme B (GzmB), and supramolecular attack particles (SMAPs), which are particles encased by sugar residues and extend cytotoxic engagement on target cells^19, 20, 21^. The secretion of these cytotoxic molecules is a tightly regulated process, with Munc13-4 being a critical priming factor for their release in CD8⁺ T cells and NK cells in both mouse and human^22, 23^. Loss-of-function mutations in Munc13-4 impair lytic granule exocytosis, leading to defective cytotoxic responses and live threatening familial haemophagocytic lymphohistiocytoses^24^. Recent evidence suggests that IFNγ, mostly recognized for its immunoregulatory functions, also plays a more direct role in cytotoxicity than previously appreciated^25^, potentially co-regulated with other cytotoxic molecules.

IFNγ is rapidly produced upon T cell activation; however, its production is independent of mature synapse formation^26, 27, 28^. It is synthesized in the endoplasmic reticulum, processed in the Golgi apparatus, and subsequently release. The secretion of IFNγ has been shown to be polarized at the synapse^1, 29, 30^, in contrast to other cytokines that are secreted multidirectionally^29^. While cytotoxic synapses restrict killing to antigenic target cells, IFNγ exerts widespread effects, highlighting its multifaceted role in immunity^30, 31^. Prolonged synapse formation sustains IFNγ transcription and production^32^. Moreover, IFNγ has been identified within SMAPs from human CD8⁺ T cells, suggesting its potential role as an effector molecule^20^. However, the composition of murine SMAPs remains unexplored, and despite these insights, the dynamics of IFNγ storage and secretion in CD8⁺ T cells, including whether it is co-packaged with other cytotoxic molecules, remain unclear.

In this study, we demonstrate that IFNγ is partially stored within GzmB⁺ cytotoxic granules in both human and murine CD8⁺ T cells. Using super-resolution imaging, we show that IFNγ exists both as a soluble molecule and encapsulated within SMAPs. Importantly, IFNγ is co-secreted with GzmB from the same granules at the immunological synapse, which we term "lytic IFNγ" due to its co-localization with cytotoxic effectors.

Moreover we identify Munc13-4 as a critical factor for early IFNγ release in murine cytotoxic T lymphocytes (CTLs). Munc13-4 deficiency impairs both lytic granule exocytosis and early IFNγ release at the immunological synapse, leading to reduced cytotoxicity in Munc13-4 knockout (KO) T cells. However, prolonged CD3 stimulation restores IFNγ secretion in Munc13-4 KO cells, suggesting the existence of alternative secretion pathways beyond synaptic release. Further imaging reveals that IFNγ is secreted at distal membrane sites during sustained synapse formation in both mouse and human CTLs. Polarization kinetics of IFNγ vesicles show that, while vesicles initially accumulate at the synapse, sustained synapse formation leads to a fraction of IFNγ vesicles being polarized to distal membrane sites. These findings indicate the presence of Munc13-4-independent release pathways for distinct populations of IFNγ. This mechanism suggests a role for IFNγ beyond cytotoxicity, potentially in immune regulation. These findings uncover a novel mechanism for IFNγ storage and release, advancing our understanding of its dual roles in cytotoxicity and immune regulation. By elucidating its dynamic release patterns and co-localization with cytotoxic molecules, this study underscores IFNγ’s critical role in CD8⁺ T cell-mediated antitumor immunity.

## Results

### IFNγ partially localizes to Granzyme B compartments in effector CTLs

To investigate the intracellular distribution of IFNγ, we first examined its production in effector CTLs following anti-CD3 stimulation. Naive murine CD8⁺ T cells were isolated from splenocytes and activated using anti-CD3/CD28 Dynabeads, resulting in the formation of blast cells after 2 days. Mouse IL-2 was added to support proliferation and maintain activation. On day 5, effector CTLs were allowed to rest for 2 hours and restimulated on anti-CD3-coated plates for 4 hours. Intracellular IFNγ production was assessed by flow cytometry (FACS). In the absence of protein secretion inhibitors, IFNγ production was rapidly induced in CTLs (Fig. 1a). Within 1 hour of restimulation, approximately 50% of the cells expressed IFNγ, increasing to over 90% by 2 hours and nearly 100% by 4 hours. Similar production kinetics were observed in both day 4 and day 5 effector CTLs (Fig. 1b).

**Figure 1:**
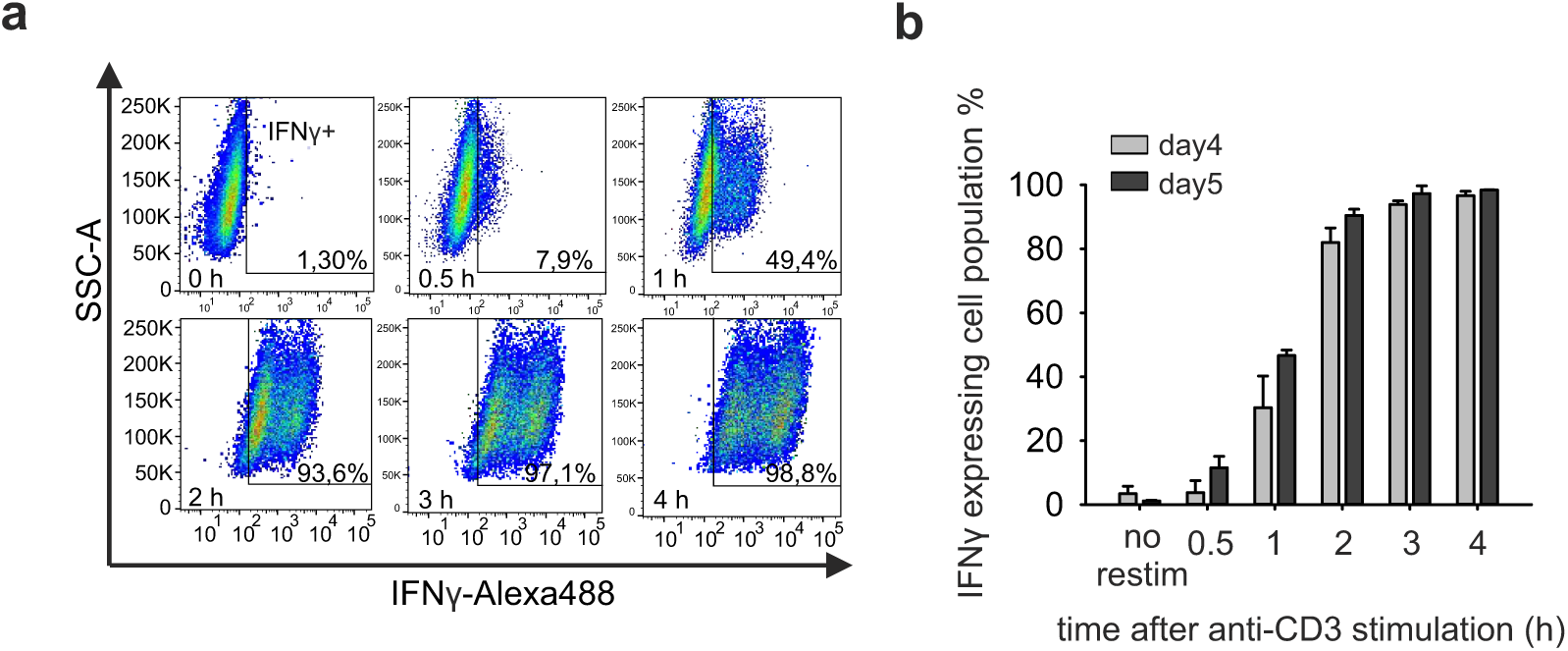
Rapid induction of IFNγ expression in mouse effector CTLs upon anti-CD3 stimulation. **a,** Day 5 effector mCTLs were stimulated on anti-CD3-coated plates, and intracellular IFNγ was stained and analyzed at the indicated time points using flow cytometry. **b,** Kinetics of IFNγ expression in day 4 and day 5 effector CTL populations, derived from analysis in (**a**). Three independent experiments were performed for FACS analysis. Data are shown as mean ± SEM. Statistical significance was determined using the Student’s t-test: *p < 0.05.

Next we analyzed the subcellular distribution of IFNγ in effector CTLs using high-resolution structured illumination microscopy (SIM). Effector cells were isolated from Granzyme B-mTFP knock-in (GzmB KI) mice^33^ in which endogenous GzmB is labeled with mTFP fluorescence and were stained with anti-IFNγ antibody. Partial co-localization of IFNγ with GzmB⁺ compartments was observed (Fig. 2a, upper panel). Similarly, overexpression of IFNγ- mCherry in GzmB KI CTLs revealed substantial co-localization of IFNγ with GzmB⁺ compartments (Fig. 2a, lower panel). Pearson’s correlation analysis confirmed partial co- localization between endogenous or overexpressed IFNγ and GzmB⁺ granules (Fig. 2b). To investigate IFNγ sorting, we overexpressed IFNγ-mCherry in day 5 effector CTLs and analyzed its localization 8 hours after transfection, at which time point IFNγ was mostly sorted out of the Golgi (Fig. 2c, upper panel). Since >90% of effector CTLs produce IFNγ within 2 hours of stimulation (Fig. 1b), we stimulated transfected cells on anti-CD3-coated plates for 2 hours and examined IFNγ distribution relative to GzmB⁺ granules (Fig. 2c, lower panel). Pearson’s correlation analysis demonstrated substantial colocalization between IFNγ and GzmB with a slight increase observed after stimulation (Fig. 2d, left). However, the number of IFNγ⁺ and GzmB⁺ vesicles does not change after 2 hours of stimulation, suggesting that newly synthesized IFNγ is targeted to pre-existing compartments without generating additional IFNγ⁺ vesicles (Fig. 2d, right).

**Figure 2:**
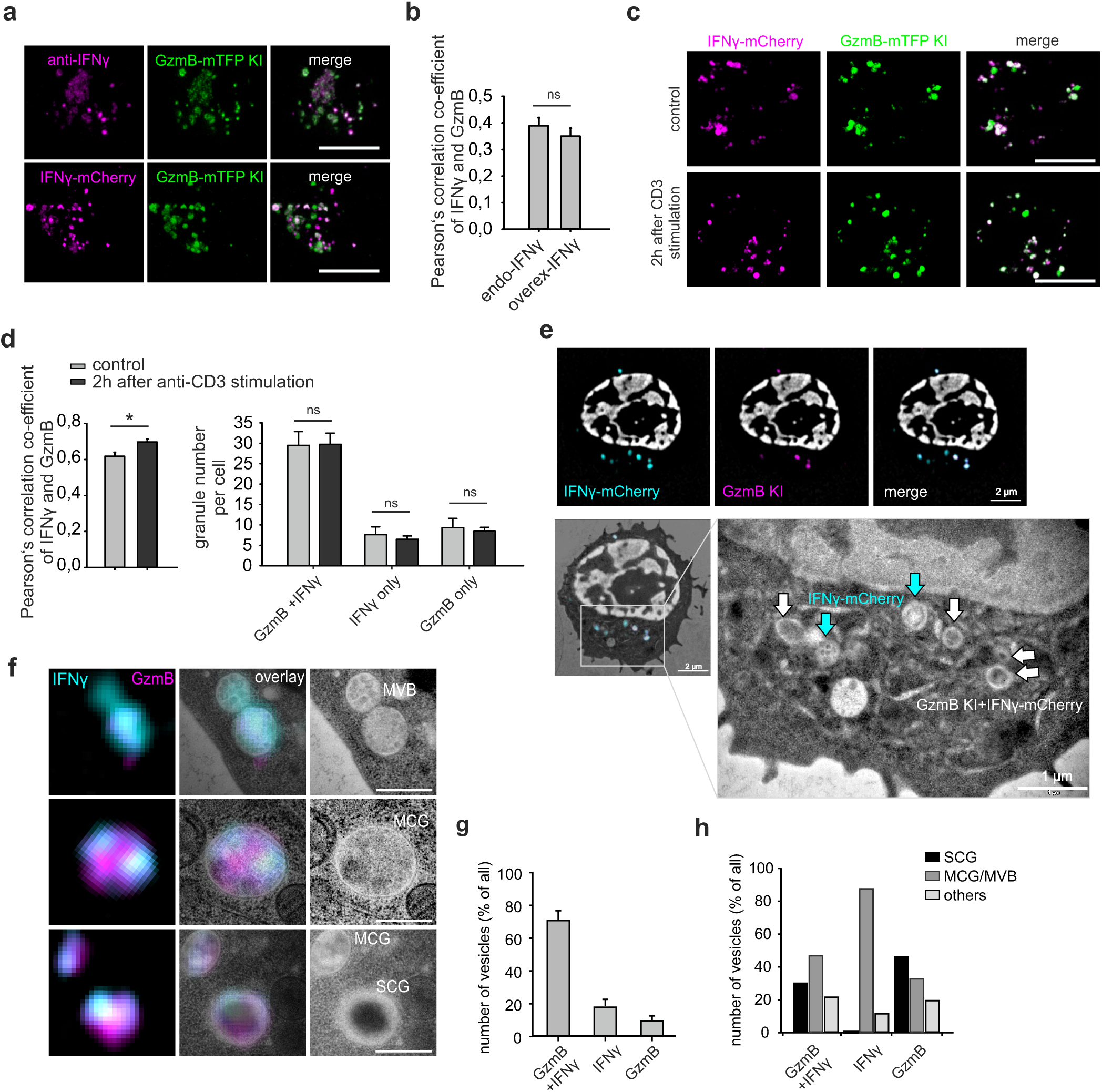
IFNγ partially localizes to Granzyme B (GzmB) compartments in mouse effector CTLs. **a,** Localization of IFNγ was analyzed in day 5 GzmB-mTFP knock-in (GzmB KI) effector CTLs using structured illumination microscopy (SIM). Endogenous IFNγ was detected with anti-IFNγ antibodies (upper panel), and overexpressed IFNγ was visualized in GzmB KI cells expressing IFNγ-mCherry (lower panel). Cells were fixed on poly-ornithine- coated plates without restimulation. Scale bar, 5 µm. **b,** Pearson’s colocalization analysis of endogenous and overexpressed IFNγ with GzmB from (**a**) (N=3 preparations; n_endo_= 85 cells for endo, n_overexp_=47 cells). **c,** SIM images of overexpressed IFNγ-mCherry in day 5 GzmB KI effector CTLs plated on anti-CD3-coated plates for 2 hours. Scale bar, 5 µm. **d,** Left: Pearson’s colocalization analysis of IFNγ-mCherry with GzmB before and after 2 hours of CD3 stimulation, derived from (**c**) Right: the number of IFNγ⁺GzmB⁺ vesicles were also quantified under the conditions described in **c** (N=3 preparations; n_control_= 17 cells for endo, n_CD3_=18 cells). **e,** Correlative light and electron microscopy (CLEM) analysis of IFNγ-mCherry-expressing day 5 GzmB KI effector CTLs. Shown are super-resolution SIM images (top row) and the overlay with the corresponding electron microscopy (EM) image (bottom panel, left). In the magnified TEM image (bottom panel, right) blue arrows point to IFNγ⁺ compartments, and white arrows denote IFNγ⁺GzmB⁺ double-positive compartments. **f,** Representative IFNγ⁺ and IFNγ⁺GzmB⁺ double-positive organelles, cropped from three different mouse CTL images as displayed in (**e**). SIM images (left), SIM/TEM overlay images (middle) and TEM images (right). Scale bar, 0.5 µm. **g,** Quantification of IFNγ⁺GzmB⁺, IFNγ⁺ and GzmB⁺ organelles identified by CLEM in percent. N=5, n_cells_=27, n_organelles_=135. **h,** Quantification of the different organelle fractions, defined by their expression profile and morphology (multicore granule, MCG; multivesicular body, MVB; single core granule, SCG and others). N=5, n_cells_=27, n_organelles_=135. Data are shown as mean ± SEM. Statistical significance was determined using the Student’s t-test: *p < 0.05.

To further investigate the ultrastructure of IFNγ⁺ compartments and their relationship to cytotoxic granules (CGs), we employed correlative light and electron microscopy (CLEM). CTLs expressing IFNγ-mCherry were high-pressure frozen, sectioned after freeze substitution, and analyzed by both SIM and transmission electron microscopy (TEM). CLEM analysis revealed diverse morphologies of IFNγ⁺ compartments. IFNγ⁺ and GzmB⁺ vesicles displayed typical CG features, including classical single-core granules (SCGs) and multicore granules (MCGs)^21^ (Fig. 2e, f). Most IFNγ⁺ compartments (75%) were double-positive for GzmB, while <20% were single-positive for either IFNγ or GzmB (Fig. 2e, g). IFNγ⁺ GzmB⁺ double-positive compartments predominantly exhibited CG structures, including SCGs and MCG/MVBs (categorized as one group), while GzmB⁺ single-positive compartments were largely SCGs. In contrast, IFNγ⁺ compartments alone exhibited multivesicular body (MCG/MVB)-like structures distinct from SCGs. Conversely, structures that did not fit these classifications were categorized as “other” (Fig. 2h).

Together, these findings indicate that IFNγ is produced upon CD3 stimulation and is partially sorted into GzmB⁺ CGs, including SCGs and MCGs.

### IFNγ in cytotoxic granules exists in both soluble and SMAP-associated forms

To determine the forms of IFNγ present in CTLs and verify its packaging in cytotoxic granules (CGs), we harvested day 5 effector CTLs from Synaptobrevin2-mRFP knock-in (Syb2 KI) mice and subjected them to nitrogen cavitation to disrupt the cells for CG isolation^34^. Syb2 is the marker of mature fusogenic CGs^35^ in mouse CTL. This marker allowed us to discriminate CGs from other organelles after sucrose density gradient centrifugation^21^. We resolved two classes of CGs: multicore granules (MCGs) in fraction 6 and single-core granules (SCGs) in fraction 8 (Fig. 3a). Isolated granules retained mRFP membrane signals and GzmB content, indicating intact structures (Fig. 3b). Immunostaining for endogenous IFNγ revealed its presence in both fraction 6 and fraction 8 granules, which also contained GzmB in day 5 effector CTLs. Quantification showed that 61.6% of MCGs and 78.1% of SCGs were positive for IFNγ (Fig. 3b). Given the presence of IFNγ in MCGs, we investigated whether it was associated with insoluble SMAPs. The majority of day 5 effector cells (69%; N=3) retained IFNγ expression following dynabeads activation without additional restimulation (Extended Data Fig. 1a,b). Co-expression of IFNγ-mCherry and TSP-1-GFPspark, a SMAP marker, in WT effector CTLs allowed us to examine IFNγ localization relative to the SMAP. High-resolution SIM revealed partial co-localization of IFNγ with TSP-1, including several small TSP-1 puncta encapsulated within IFNγ⁺ compartments (Fig. 3c, white arrows). Manders’ colocalization analysis confirmed greater colocalization of TSP-1 with IFNγ than the reverse, consistent with TSP-1 encapsulation within IFNγ+ compartments (Fig. 3d).

**Figure 3:**
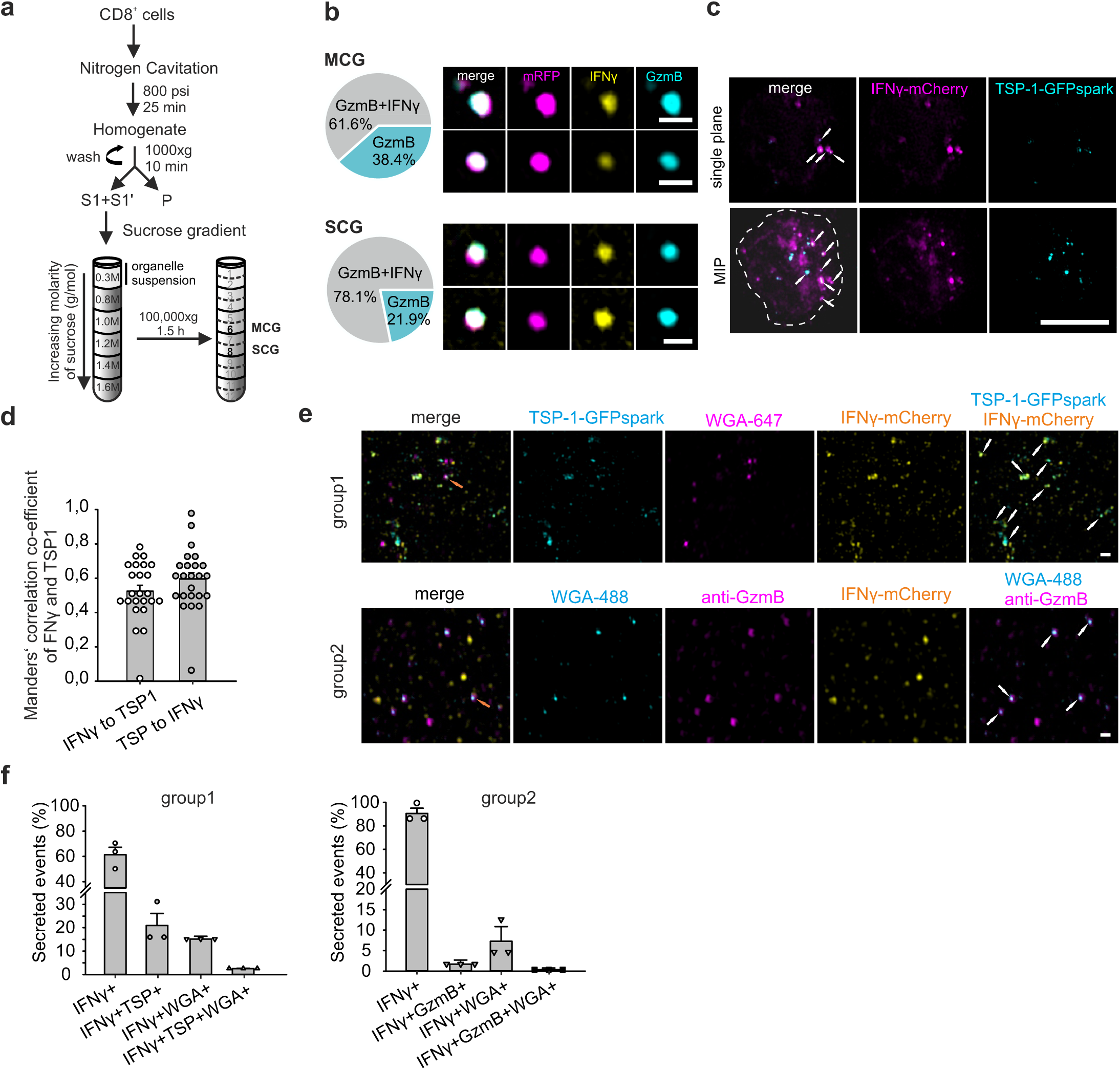
IFNγ in cytotoxic granules (CGs) is present in both soluble and SMAP-associated forms. **a,** Schematic representation of the CG isolation procedure using discontinuous sucrose density gradient ultracentrifugation. Sucrose fractions enriched for multi-core granules (MCGs; fraction 6) and single-core granules (SCGs; fraction 8) generated from Syb2-mRFT knock-in effector T-cells were collected. **b,** Left: Fraction of IFNγ⁺GzmB⁺ organelles within MCGs and SCGs (N_MCG_=3, n_MCG_=323, N_SC_=2, n_SCG_=32). Right: SIM images of representative MCGs and SCGs isolated from the respective fractions, stained with anti- IFNγ (yellow) and anti-GzmB-Alexa647 (cyan) antibodies. Syb2-mRFP (magenta) labels intact organelles. Scale bar, 0.5 µm. **c,** SIM images of day 5 WT effector T cells co-expressing IFNγ- mCherry and TSP-GFPspark. Single-plane and maximum intensity projection (MIP) images are presented. The white dotted line outlines the cell footprint. White arrows indicate regions of IFNγ and TSP-1 co-localization. Scale bar, 5 µm. **d,** Manders’ colocalization analysis of IFNγ-mCherry and TSP-GFPspark signals from (**c**) (N=3 preparations; n=24 cells). **e,** Day 5 effector T cells transfected with either IFNγ-mCherry and TSP-1-GFPspark (upper panel) or IFNγ-mCherry alone (lower panel) were plated on a supported bilayer coated with anti-CD3 to induce synapse formation. After 90 minutes of incubation at 37°C, secreted particles were analyzed for colocalization with SMAP markers, including granzyme B and WGA. White arrows indicate double-positive particles, and orange arrows indicate triple-positive particles. Scale bar, 0.5 µm. **f,** Quantification of single-, double-, and triple-positive particles from (**e**) presented as percentages (N=3 preparations; n_group1_=45 fields of view, with 18569 particles analyzed; n_group2_=41 fields of view, with more than 79994 particles analyzed). Data are presented as mean ± SEM, with the distribution of individual data points shown.

To further investigate whether IFNγ is associated with secreted SMAPs, we analyzed its co- localization with SMAP markers, including WGA-binding glycoprotein, TSP-1, and GzmB, on the secreted particles. Two groups of day5 effector CTLs were prepared: one co-expressing IFNγ-mCherry and TSP-1-GFPspark, and another expressing IFNγ-mCherry alone. Cells from both groups were stimulated for 90 minutes on supported lipid bilayers coated with ICAM-1 and anti-CD3 to induce synapse formation and granule secretion. Post-stimulation, cells were washed with D-PBS, leaving secreted materials deposited on the bilayers. Secreted materials were stained with WGA to label sialic acid residues enriched on SMAPs, and the second group was additionally stained with anti-GzmB to identify cytotoxic SMAPs (Fig. 3e). SIM imaging revealed a heterogeneous population of secreted vesicles and particles on the bilayers.

Numerous IFNγ single positive events were observed, with subsets of IFNγ co-localizing separately with TSP-1, WGA, or GzmB as double-positive events. Object-based colocalization analysis confirmed that a fraction of secreted IFNγ co-localized with each of the tested SMAP markers (Fig. 3f). Due to the overexpression of IFNγ-mCherry, large IFNγ-only puncta were observed that were not associated with the SMAP markers as TSP-1, WGA or GzmB. However, these IFNγ puncta might exist as protein aggregates containing IFNγ, which appear as visible fluorescent dots, or be associated with exosomes or SMAP-like particles, which feature different markers or other effector molecules. These results demonstrate that IFNγ exists in cytotoxic granules both in soluble form and SMAP-associated form, underlining its diverse modes of packaging within effector CTLs.

### IFNγ and Granzyme B co-release at the immune synapse in mouse effector CTLs

We next investigated the real-time dynamics of IFNγ release from cytotoxic granules (CGs) using confocal imaging and total internal reflection fluorescence microscopy (TIRFM), which offers high axial resolution and enables the visualization of individual granules in T cells. IFNγ- mCherry was expressed in Granzyme B (GzmB)-mTFP knock-in (KI) effector T cells to monitor IFNγ polarization upon target cell engagement. Live-cell confocal imaging revealed that IFNγ and GzmB polarized together toward the immunological synapse within 3 minutes of conjugation with P815 target cells (Fig. 4a). For TIRFM, T cells were stimulated on anti-CD3- coated glass coverslips to form synapses, with additional extracellular calcium added to trigger granule release. Two distinct release patterns were observed: (1) IFNγ release in a soluble form or (2) IFNγ release accompanied by detectable particle deposition at the synapse. Soluble release was characteristic of single-core granules (SCGs), whereas deposition of particles was associated with multicore granules (MCGs) (Fig. 4b). Individual vesicle release events were visualized and analyzed. SCG release events showed full collapse of the vesicle at the synaptic membrane, marked by a sudden disappearance of diffusive IFNγ-mCherry and GzmB-mTFP fluorescence (Fig. 4b, upper panel). Fluorescence intensity profiles confirmed a rapid loss of signal at the time frame when the fusion pore opened (indicated by an asterisk; Fig. 4c, upper panel). In contrast, MCG release events resulted in particle deposition after the release of soluble IFNγ and GzmB from the same vesicle (Fig. 4b, lower panel). The deposited IFN-γ and GzmB signals persisted after the soluble phase dissipated, as evidenced by the elevated baseline fluorescence intensity following the fusion event (Fig. 4c, lower panel). Similar to SCGs, the majority of the IFNγ signal in MCGs rapidly diffuses away, highlighting the fact that both soluble IFNγ and SMAP-encapsulated IFNγ are present in MCGs (Fig. 4c, lower panel). Quantitative analysis revealed that the majority of release events (93.2%) corresponded to SCGs, with only 6.8% representing MCGs (Fig. 4d). Among the released granules, 60.9% were positive for both IFNγ and GzmB, while 22.6% contained only GzmB and 16.5% contained only IFNγ (Fig. 4e, left). The composition of deposited SMAPs at the synapse showed a similar distribution to that of the granule contents: the majority encapsulated both IFNγ and GzmB (66.6%), while 23.8% contained only GzmB and 9.5% contained only IFNγ (Fig. 4e, right). These findings highlight that the majority of IFNγ release at the synapse occurs in conjunction with GzmB, both in soluble and SMAPs-associated form. We term this population of IFNγ released alongside cytotoxic mediators “lytic IFNγ”.

**Figure 4:**
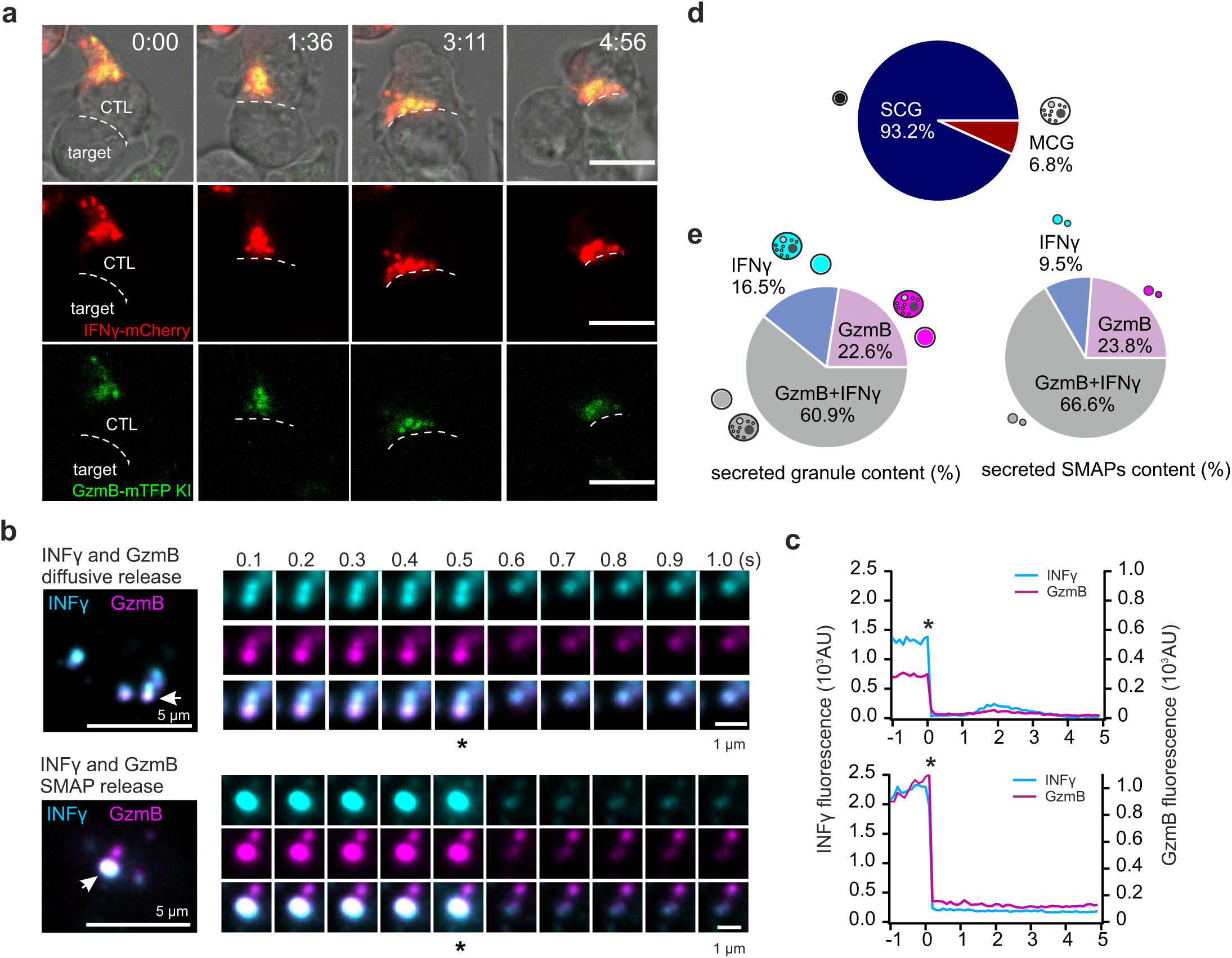
IFNγ and Granzyme B co-release at the immune synapse in mouse effector CTLs. **a,** Live-cell confocal images of an IFNγ-mCherry-expressing GzmB KI CTL forming a synapse with a P815 target cell. Scale bar, 5 µm. **b,** Day 5 IFNγ-mCherry-expressing GzmB KI CTL plated on anti-CD3-coated glass coverslips to induce granule secretion. Cells were perfused with 10 mM extracellular calcium buffer, and secretion events were recorded by TIRFM. Snapshots show secreting granules (white arrows) at the synapse in individual cells (left panel); asterisks indicate the time frame when the fusion pore opens (right panels). Two secretion patterns were shown: IFNγ⁺GzmB⁺ double-positive granules with SMAPs (top) and diffusive release of IFNγ⁺GzmB⁺ granules (bottom). Images were acquired at 10 Hz. Scale bar, 1 µm. **c,** Fluorescence intensity profiles of granule release from (**b**) over 5 s. A sudden decrease in IFNγ and GzmB fluorescence indicates fusion events. **d,** Classification of granule secretion at the synapse. Granules exhibiting IFNγ⁺GzmB⁺ co-release, IFNγ-only and GzmB- only release were categorized based on SMAP deposition (MCG) or diffusive release (SCG) (N=4 preparations; n = 274 cells). **e,** Quantification of granules categorized by their secreted content of GzmB and/or IFNγ at the synapse (N=4 preparations; n = 381 cells). Left, percentages of double-positive or single-positive granule secretion events. Right, proportion of secreted GzmB and IFNγ particles (N=2 preparations; n_cell_= 10, n_smaps_=21).

### IFNγ partially localizes to Granzyme B compartments in human effector CTLs

To determine whether the intracellular localization of IFNγ within cytotoxic granules (CGs) observed in mouse effector T cells is conserved in humans, we performed colocalization studies in human CTLs. CD8⁺ T cells were isolated from peripheral blood mononuclear cells (PBMCs), stimulated with anti-CD3/CD28-coated Dynabeads, and were allowed to proliferate for 5 days. A fraction of the day 5 human effector CTL population retained IFNγ expression, with 6.7% IFNγ⁺ cells detected by flow cytometry (Extended Data Fig. 2a,b). These effector CTLs were stained for endogenous IFNγ and GzmB. SIM imaging revealed partial co- localization of IFNγ with GzmB in these cells (Fig. 5a). Endogenous IFNγ appeared as small puncta, with occasional larger vesicular structures on poly-L-ornithine-coated coverslips under unstimulated conditions (Fig. 5a, upper panel). Upon brief (10 min) anti-CD3 stimulation, larger IFNγ puncta increased as synapses formed (Fig. 5a, lower panel). Pearson’s and Manders’ co-localization analyses confirmed partial co-localization of IFNγ with GzmB, with Manders’ analysis indicating a higher co-localization value for IFNγ with GzmB than for GzmB with IFNγ, suggesting the presence of two IFNγ compartments: one overlapping with GzmB and the other independent of it (Fig. 5c,d). Notably, short-term CD3 stimulation did not significantly alter colocalization patterns. Overexpression of hIFNγ-mNeonGreen and hGzmB-mCherry in effector CTLs resulted in a high co-localization pattern (Fig. 5b). Quantification revealed that over 90% of granules were double-positive, containing both IFNγ and GzmB (Fig. 5e). This result is consistent with the high Pearson’s correlation values observed (Fig. 5f). The remaining fraction consisted of single-positive granules, containing either IFNγ or GzmB, supporting the existence of two distinct IFNγ populations in human CTLs.

**Figure 5:**
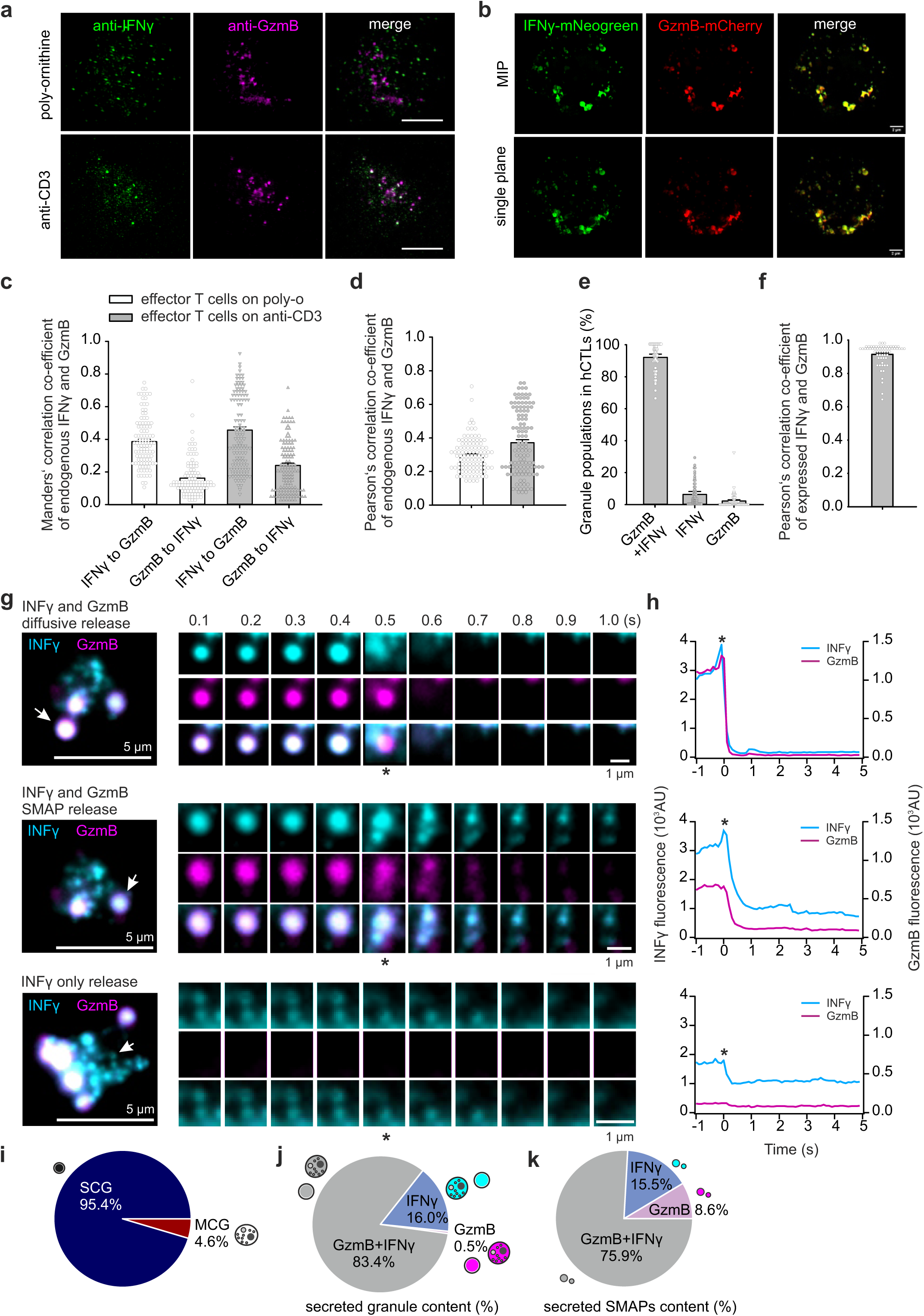
IFNγ partially localizes to Granzyme B (GzmB) compartments in human effector CTLs. **a,** SIM images of day 5 human effector CTLs with endogenous IFNγ and GzmB stained using specific antibodies. Cells were fixed on poly-ornithine- or anti-CD3-coated glass coverslips after 10 min. Scale bar, 2 µm. **b,** SIM images of CTLs overexpressing IFNγ- mNeonGreen and GzmB-mCherry. Cells were fixed on poly-ornithine-coated coverslips without restimulation. Scale bar, 2 µm. **c,** Manders’ colocalization and **d,** Pearson’s colocalization analysis of IFNγ-mNeonGreen and GzmB-mCherry signals from (**a**) (N=3 donors; n = 105 cells). **e,** Pearson’s colocalization analysis of IFNγ-mNeonGreen and GzmB- mCherry signals from (**b**) (N=3 donors; n = 51 cells). **f,** Quantification of object-based colocalization of IFNγ-mNeonGreen and GzmB-mCherry granules (N=3 donors; n_cells_ = 51; n_granules_= 63). Data are presented as mean ± SEM, with the distribution of individual data points shown. **g,** Day 5 CTLs transfected with IFNγ-mNeonGreen and GzmB-mCherry were plated on anti-CD3-coated glass coverslips to induce granule secretion for TIRFM analysis. Three secretion patterns were observed: IFNγ⁺GzmB⁺ double-positive granules with SMAPs (top), diffusive release of IFNγ⁺GzmB⁺ granules (middle), and IFNγ-only release (bottom). Snapshots show secreted granules (white arrows) at the synapse of individual cells (left panel); asterisks indicate the time frame when the fusion pore opens. Images were acquired at 10 Hz. Scale bar, 1 µm. **h,** Fluorescence intensity profiles of granule release from (**g**) over 5 s. A sudden decrease in IFNγ and GzmB fluorescence indicates fusion events. **i**, Characterization of CG secretion at the synapse. Granules exhibiting IFNγ⁺GzmB⁺ co-release, IFNγ only and GzmB-only release were categorized based on SMAP deposition (MCG) or diffusive release (SCG) (N=3 donors; n = 76 cells). **j,** Quantification of GzmB and IFNγ secretion at the synapse. Percentages of double-positive or single-positive granule secretion events (N=3 donors; n_cells_= 76; n_fusion events_= 298). **k**, Proportions of secreted GzmB and IFNγ particles (N=3 donors; n_cells_= 57; n_smaps_= 98).

To investigate IFNγ secretion at the immune synapse, we monitored individual granule release by TIRFM. Human CTLs transfected with hIFNγ-mNeonGreen and hGzmB-mCherry were plated on anti-CD3-coated coverslips to induce synapse formation, with additional extracellular calcium applied to enhance secretion probability. The majority of secreted IFNγ originated from GzmB⁺ granules, with a diffuse release pattern characterized by a sudden loss of fluorescence for both mNeonGreen and mCherry, indicative of soluble secretion (Fig. 5g,h; upper panel). A fraction of IFNγ was packaged in MCGs and co-released with GzmB, leaving behind insoluble SMAP particles at the synapse. These SMAPs retained minimal fluorescence, reflecting their deposition (Fig. 5g,h; middle panel). Additionally, a distinct population of small IFNγ-only vesicles polarized to the synapse and fused upon CD3 stimulation, releasing soluble IFNγ without GzmB. This population, termed non-lytic IFNγ, is smaller than the population of lytic granules and lacks lytic components such as GzmB (Fig. 5g,h; lower panel). Secretion analyses in human CTLs closely mirrored findings in mouse CTLs. SCGs accounted for the majority of fusion events (93.2%), with MCGs comprising the remainder (6.8%) (Fig. 5i). The majority of IFNγ fusion events were associated with GzmB (lytic IFNγ, 83.4%), while IFNγ-only fusion events accounted for 16% (Fig. 5j). Among secreted SMAPs, most contained both IFNγ and GzmB (75.9%), with fewer particles containing only IFNγ (15.5%) or only GzmB (8.6%).

Together, these results demonstrate the conservation of lytic IFNγ in human effector CTLs and its distinction from non-lytic IFNγ. Detailed fusion analyses reveal a high degree of similarity between human and mouse CTLs in the packaging and release dynamics of IFNγ and GzmB.

### Impaired lytic IFNγ release is accompanied by reduced cytotoxicity in mouse CTLs

To investigate the molecular mechanisms underlying IFNγ release, we focused on Munc13-4, a priming factor essential for lytic granule exocytosis in CTLs^36^. CTLs from Munc13-4 knockout mice (Jinx mouse^37^), in which the mutated Munc13-4 gene produces a nonfunctional protein, were compared to WT cells. Day 5 effector WT and KO cells showed similar IFNγ mRNA expression levels (Extended Data Fig. 3a) and followed a comparable upregulation of IFNγ gene expression profile upon anti-CD3 stimulation. IFNγ reached maximal expression after 2 hours of stimulation (Extended Data Fig. 3b), with no significant differences observed between WT and KO cells. To assess IFNγ secretion, we stimulated WT and Munc13-4 KO effector CTLs on anti-CD3-coated plates and performed a time-course analysis over a 20-hour period. IFNγ secretion was significantly impaired in KO cells during the first hour of stimulation. A flow cytometry-based LEGENDplex assay revealed that during the first 60 minutes of CD3 stimulation, WT T cells produced measurable levels of IFNγ (up to 70 pg/ml), whereas KO T cells exhibited a severe impairment, resulting in minimal IFNγ concentrations (up to 10 pg/ml) in the supernatant (Fig. 6a). However, after 2 hours of stimulation, ELISA assays showed that IFNγ concentrations in both WT and KO cultures reached comparable levels (around 62,000 pg/ml), indicating that KO T cells restored IFNγ secretion during sustained stimulation (Fig. 6b). Constitutive IFNγ release in KO cells exhibited a slight but not significant increase in secretion in the absence of stimulation (Extended Data Fig. 3c).

**Figure 6:**
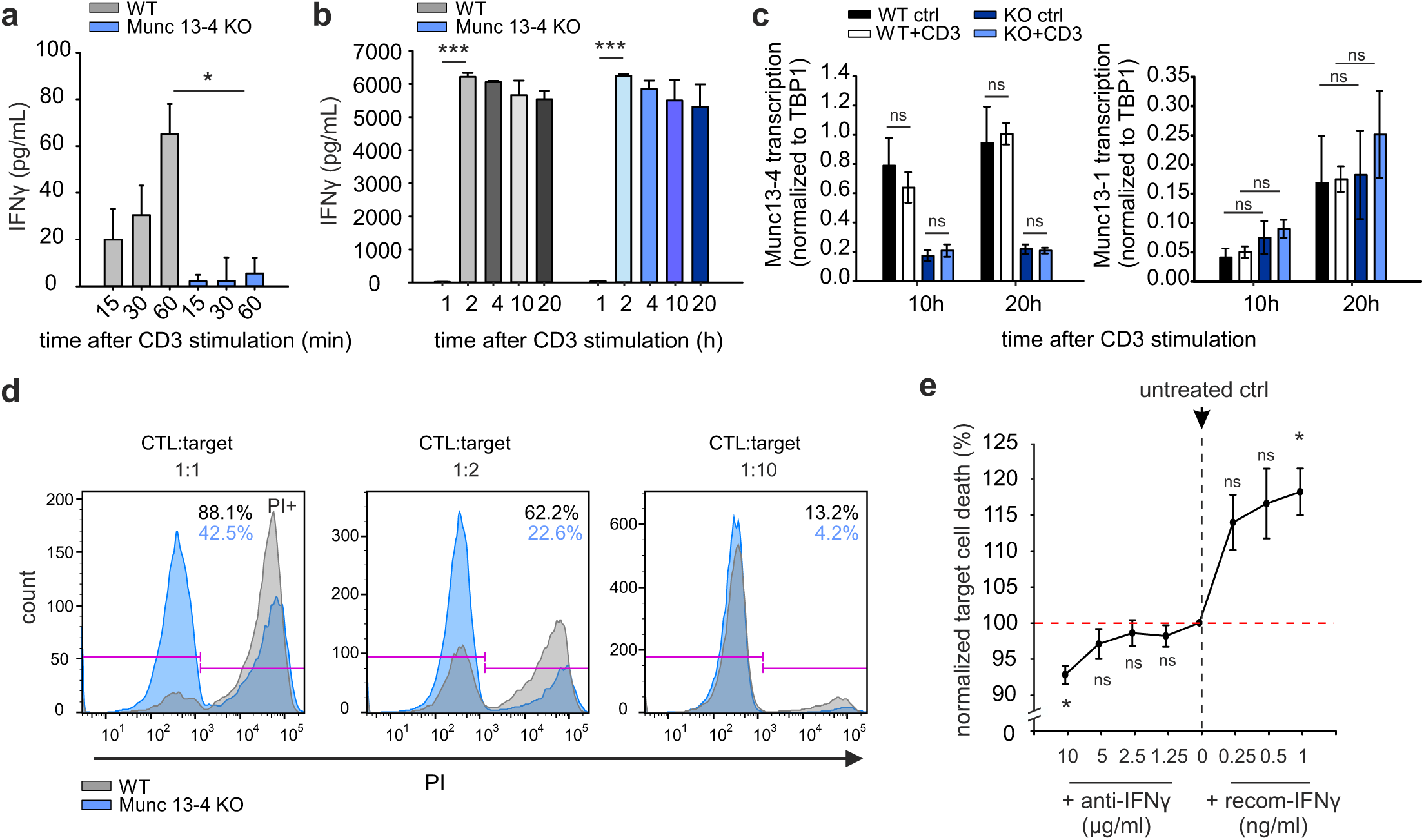
Impaired lytic IFNγ release leads to reduced cytotoxicity in mCTLs. **a, b**, IFNγ release was measured in the supernatant using a LEGENDplex kit within 60 minutes (**a**) or an ELISA kit (**b**) within 1–20 hours. Wild-type (WT, gray) and Munc13-4 knockout (KO, blue) day 5 effector CTLs were plated on anti-CD3-coated plates. Supernatants were collected at the indicated time points (N=3 for LEGENDplex analysis; N=3 for ELISA analysis). **c,** Normalized Munc13-1 and Munc13-4 mRNA transcription in WT and Munc13-4 KO CTLs at 10 h and 20 h after anti-CD3 stimulation (N=3). **d,** FACS-based killing assay of WT (grey) and Munc13-4 KO (blue) T cells. CTLs were co-cultured with p815 target cells at the indicated T cell-to-target ratios for 3 h. Dead cells were stained with propidium iodide (PI) to evaluate cell death. One representative experiment from 4 independent preparations. **e,** Effect of anti-IFNγ antibody neutralization (left quadrants) or additional recombinant IFNγ protein (right quadrants) on CTL-mediated killing. Day4 effector WT CTLs were mixed with p815 target cells as in (**d**). Dead cells were stained with PI. Untreated control cells are shown at the midpoint of the curve (N=3). Data are presented as mean ± SEM. Statistical significance was determined using the Student’s t-test: *p < 0.05, *p < 0.001; ns indicates non-significance.

Notably, overexpression of IFNγ-mCherry in effector CTLs resulted in high levels of constitutive secretion, even in the absence of CD3 stimulation in WT cells (Extended Data Fig. 4). This suggests that IFNγ release can occur independently of stimulation when protein expression levels are elevated, implying the presence of an additional IFNγ vesicle population distinct from the lytic IFNγ granule population. Since Munc13-4 KO cells exhibit normal IFNγ mRNA expression, the restoration of IFNγ release after 2 hours of stimulation indicates the involvement of a compensatory, Munc13-4-independent mechanism during prolonged activation. Munc13-1, another vesicle priming factor expressed in mouse CTLs, shares functional similarities with Munc13-4 and is a candidate for this compensatory role^36^. To explore this, we analyzed munc13-1 expression in WT and KO CTLs at 10 and 20 hours after stimulation. qPCR analysis showed no significant differences in munc13-1 expression between genotypes (Fig. 6c), suggesting that late-phase IFNγ secretion arises from non-synaptic sources rather than compensatory upregulation of Munc13-1. Furthermore, our TIRFM imaging revealed that cytotoxic granule secretion occurs predominantly during early stimulation, within 10 minutes of synapse formation. Subsequently, granule secretion is rarely detected, despite the majority of cytotoxic granules remaining polarized at the synapse (Fig. 4 and Fig. 5). This early-phase IFNγ release appears to be crucial for target cell cytotoxicity. Although granzymes and perforin are known to be the major contributors to granule-mediated killing, the fact that IFNγ co-releases with GzmB and concentrates at the synaptic cleft suggests that IFNγ may also contribute directly or indirectly to cytotoxicity. Munc13-4 KO CTLs, which are deficient in CG secretion affects early IFNγ secretion, exhibited reduced killing of P815 target cells in 2-hour co-culture assays (Fig. 6d). Neutralizing antibodies against IFNγ further decreased killing efficiency in WT CTLs, while recombinant IFNγ enhanced cytotoxicity in WT cells, demonstrating that suspended IFNγ contributes as an effector molecule in target cell killing (Fig. 6e). Notably, recombinant IFNγ protein alone does not induce cell death, suggesting that IFNγ requires cytotoxic molecules such as granzyme B to synergize its cytotoxic effects (Extended Data Fig. 3d). These findings emphasize the importance of Munc13-4-dependent synaptic secretion of IFNγ during early stimulation, which might be essential for effective target cell killing. While compensatory mechanisms restore late-phase secretion in the absence of Munc13-4, they cannot compensate for the impaired cytotoxicity. This highlights the distinct roles of IFNγ, possibly differentiating between lytic and non-lytic IFNγ populations.

### IFNγ secretion at the distal plasma membrane in mouse and human CTLs

To distinguish between lytic and non-lytic IFNγ populations, we analyzed the secretion site and distribution of IFNγ in day 5 effector mouse and human CTLs. WT and Munc13-4 KO mouse CTLs transfected with IFNγ-mCherry were stimulated on anti-CD3-coated coverslips to form synapses and secrete IFNγ. The tight synapse formation on the glass surface restricted cell migration and defined the synaptic area, where secreted materials were retained within the sealed synaptic cleft. IFNγ⁺ particles were detected outside the cell area, most likely secreted from the distal plasma membrane (Fig. 7a). While the majority of soluble IFNγ diffused into the supernatant and could not be visualized, we detected a minority of particles carrying IFNγ, likely exosome-associated, which may have originated from the same secretion site at the distal membrane. In mouse CTLs, secreted IFNγ⁺ particles were observed in both WT and Munc13-4 KO cells within 2 hours, with no significant difference in the number of secreted particles, supporting the ELISA data indicating that KO cells release IFNγ at the distal membrane despite the defect in CG exocytosis at the synapse. Quantification of these particles in cell-free regions on the coverslip revealed a modest increase in secretion over 2 hours of stimulation in WT cells. Munc13-4 KO cells also showed a trend toward increased particle release, although this was not statistically significant (Fig. 7b). It is important to note a discrepancy between the detected particles on the coverslip and the measured IFNγ concentration in the supernatant. These secreted particles were present in very low numbers on the coverslip and may not fully represent the soluble IFNγ measured in the supernatant. Therefore, this data emphasizes the fact that IFNγ is secreted at the distal membrane rather than providing a quantitatively accurate measure of released IFNγ. Similarly, distal secretion of IFNγ was observed in IFNγ-mNeogreen-transfected human CTLs. Due to the low transfection efficiency, particles were counted within a 30 µm radius of transfected cells. We observed an average of 2-4 particles per cell within 2 hours of anti-CD3 stimulation (Fig. 7c,d). These findings demonstrate that IFNγ secretion occurs at the distal plasma membrane upon anti-CD3 stimulation in both mouse and human CTLs.

**Figure 7:**
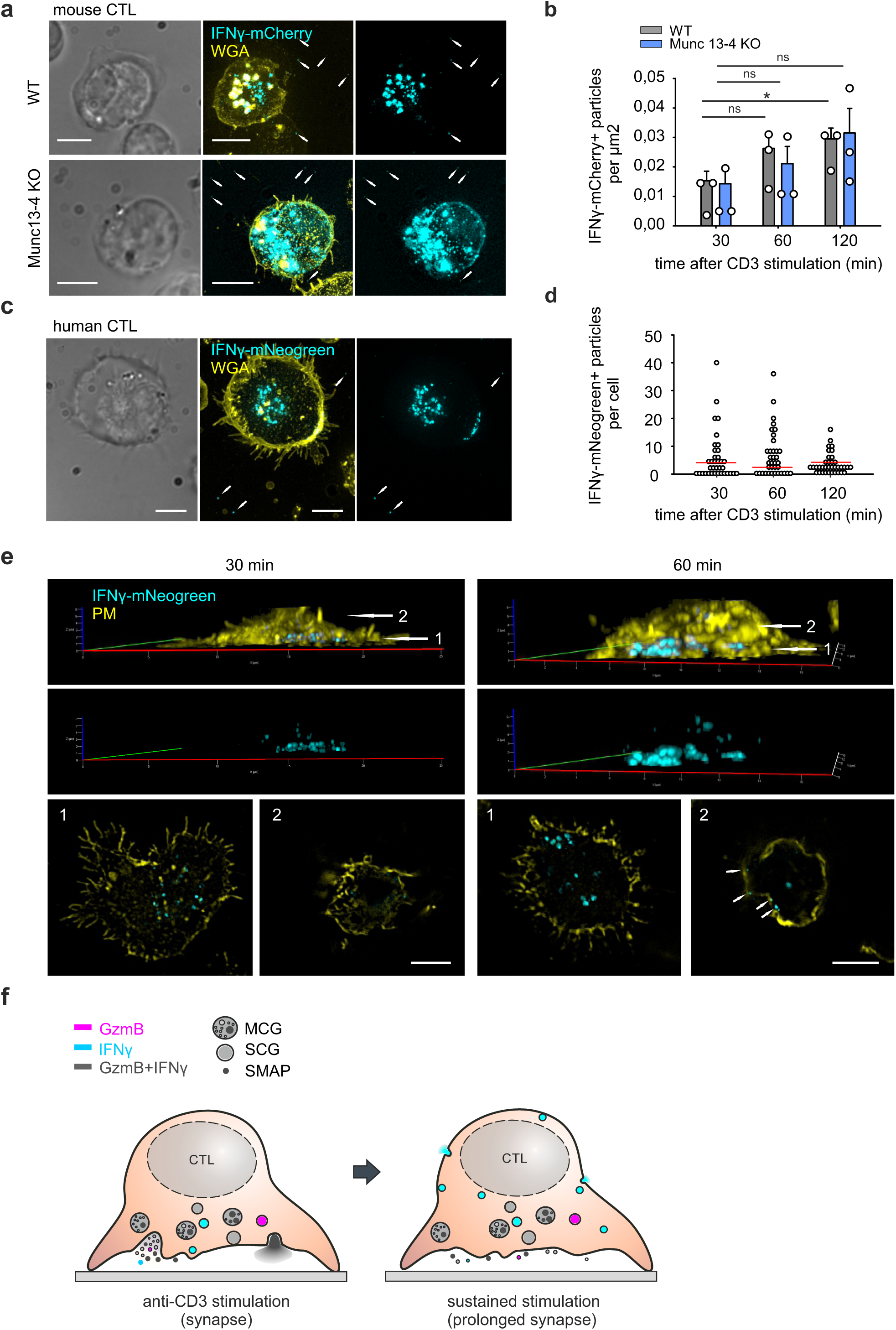
IFNγ secretion at the distal plasma membrane in mouse and human CTLs. **a**, Representative SIM images of day 5 effector WT and Munc13-4 KO mouse CTLs fixed after 60 min on anti-CD3-coated coverslips. CTLs were transfected with IFNγ-mCherry (cyan) and stained with WGA (yellow) to mark the plasma membrane. White arrows indicate secreted IFNγ-mCherry⁺ particles outside the cells. Scale bar, 5 μm. **b,** Quantification of secreted IFNγ- mCherry⁺ particles in the extracellular regions from (**a**) (N = 3 mice; n = 25-30 areas per group). **c,** Representative SIM images of day 5 effector human CTLs fixed after 60 min on anti-CD3- coated coverslips. CTLs were transfected with IFNγ-mNeonGreen (cyan) and stained with WGA (yellow) to mark the plasma membrane. White arrows indicate secreted IFNγ- mNeonGreen⁺ particles outside the cells. Scale bar, 5 μm. **d,** Quantification of secreted IFNγ- mNeonGreen⁺ particles from (**c**) at indicated time points (N = 3 donors; n = 36 cells per time point). **e,** Day 5 effector human CTLs transfected with IFNγ-mNeonGreen were fixed after 30 min (left) and 60 min (right) on anti-CD3-coated coverslips. Plasma membranes were stained with WGA. Upper panels show 3D reconstructions. Two distinct z-positions namely synaptic (1) and distal (2) are defined by arrows. Lower panels show corresponding single-plane xy images. White arrows in the distal layer (position 2) at 60 min indicate IFNγ⁺ vesicles attached to the plasma membrane away from the synapse. Scale bar, 5 μm. **f,** Schematic illustration of IFNγ secretion in CTLs. Lytic IFNγ is packaged within cytotoxic granules containing granzyme B (GzmB), including single-core granules (SCGs) and multi-core granules (MCGs), and is also associated with SMAPs. IFNγ is predominantly co-released with GzmB at immune synapses (left), with prolonged synapse formation also leading to IFNγ release at distal plasma membrane sites (right).

As IFNγ is predominantly polarized and secreted at the synapse, we next investigated whether IFNγ⁺ vesicles also polarize to the distal membrane. IFNγ-expressing CTLs were fixed on anti- CD3-coated coverslips at 30 and 60 minutes post-stimulation. 3D SIM imaging revealed an accumulation of IFNγ at the synapse within 30 minutes as expected (Fig. 7e, left; Extended Data Fig. 5a, left). However, sustained stimulation led to a fraction of IFNγ⁺ vesicles polarized to the distal plasma membrane. Single-plane 2D images confirmed predominant granule accumulation at the synapse (position 1, synaptic layer) during early stimulation, with minimal localization to the cytosol (position 2, distal membrane layer). By 60 minutes, synaptic IFNγ pool had decreased, while a fraction of IFNγ⁺ vesicles appeared to polarize to the distal membrane in both mouse and human CTLs (white arrows, position 2, lower panel), a feature that was absent during earlier time points (Fig. 7e, right; Extended Data Fig. 5a, right). These vesicles likely represent a non-lytic IFNγ population primed for secretion during sustained synaptic activation.

These findings reveal that IFNγ is secreted not only at the immune synapse but also at the distal plasma membrane, potentially driven by sustained synapse formation in mouse and human CTLs. Taken together, our study uncovers a previously unrecognized mechanism of IFNγ packaging within GzmB⁺ cytotoxic granules, termed ’lytic- IFNγ.’ Lytic-IFNγ is released at the synapse in both soluble and SMAP-associated forms, contributing to the cytotoxicity of CTLs. Furthermore, sustained synaptic activation potentially drives the multidirectional secretion of IFNγ at the distal membrane (Fig. 7f).

## Discussion

Effector CD8⁺ T cells play essential roles in antitumor immunity. Beyond their cytotoxic functions, CD8⁺ T cells secrete IFNγ to modulate immune responses and enhance antitumor activity^9, 11, 14, 31, 38^. Despite its critical role in antitumor immunity, the direct contribution of IFNγ to cytotoxicity remains unclear. In this study, we identify a fraction of IFNγ stored within GzmB⁺ cytotoxic granules, termed "lytic IFNγ". This finding redefines IFNγ as an integral component of the granule-mediated killing machinery, acting in concert with GzmB. Using advanced imaging techniques, we demonstrate that lytic IFNγ is released in both soluble and supramolecular attack particle (SMAP)-associated forms, co-released with GzmB at the immune synapse in human and mouse CD8+ T cells (Figs. 4 and 5). Using Munc13-4-deficient mouse T cells, we identified an intriguing phenotype where IFNγ release was impaired within the first hour but returned to normal after 2 hours of sustained synaptic engagement (Fig. 6a,b). While TIRF imaging revealed no GzmB⁺ vesicle secretion at the synapse in Munc13-4 KO T cells, this suggests that the later released IFNγ may originate from the distal membrane rather than the synaptic cleft.

GzmB secretion is restricted to the synaptic cleft, ensuring specificity in target cell killing. Similarly, co-released synaptic “lytic IFNγ” is concentrated at the cleft, suggesting a potentially distinct functional role compared to IFNγ released at the distal membrane. IFNγ plays a pivotal role in CD8⁺ T cell-mediated tumor cytotoxicity by enhancing immune recognition and destruction of tumor cells. It upregulates MHC class I molecules on tumor cells, improving antigen presentation and making tumor cells more visible to CD8⁺ T cells^39, 40^. Furthermore, IFNγ sensitizes tumor cells to apoptosis by increasing the expression of pro-apoptotic caspases. It also enhances the cytolytic activity of CD8⁺ T cells by promoting the expression of perforin, granzymes, and Fas ligand, which are critical for inducing tumor cell apoptosis^41, 42^. We speculate that synaptic lytic IFNγ contributes to the acute reprogramming of target cells, sensitizing them to T cell-mediated attack by creating a concentrated microdomain of immune signaling at the synapse, if not directly exerting cytotoxic effects. To investigate this, we applied an anti-IFNγ antibody before T cells and target cells conjugated to neutralize both suspended IFNγ and synaptic IFNγ. This neutralization reduced T cell cytotoxicity. In contrast, the addition of recombinant IFNγ (1 ng/ml) slightly enhanced target cell killing (Fig. 6e). This extracellular IFNγ likely sensitizes tumor cells, enhancing killing efficiency in addition to granule-mediated death. Notably, recombinant IFNγ alone without T cells did not induce target cell death (Extended Data Fig. 3d), suggesting that IFNγ requires co-released cytotoxic molecules to synergize cytotoxic responses.

The expression of effector molecules in T cells is highly adaptable to environmental cues and compensatory needs^32, 43, 44^. For instance, although IFNγ expression correlates with cytotoxicity, IFNγ-negative CD4⁺ and CD8⁺ T cells can still produce other immune mediators, such as perforin, TNFα, and IL-2, in response to viral antigens, thereby maintaining their effector functions^45^. Interestingly, overexpression of IFNγ in CD8⁺ T cells alters granule secretion patterns, favoring the release of single-core granules (SCGs) over multicore granules (MCGs) (Figs. 4d and 5i), a pattern distinct from what we previously observed in day 5 effector mouse T cells^21^. This finding suggests that T cells may adapt their granule secretion patterns in response to the local immune environment, optimizing the release of effector molecules to enhance their activity/effector function under conditions of massive IFNγ production (i.e. acute inflammation).

Our findings also reveal structural heterogeneity within cytotoxic granules (CGs), with IFNγ distributed across both SCGs and MCGs (Figs. 2e,f and 3). MCGs appear to serve as reservoirs for diverse contents, including soluble molecules, exosomes, and SMAPs^21^. Secreted particles from CD8⁺ T cells exhibit remarkable diversity, including double-positive particles containing both IFNγ and GzmB, as well as single-positive particles with either IFNγ or GzmB (Figs. 4 and 5). These IFNγ-containing particles associated with various SMAP markers (Figs. 3e,f) suggest a dynamic sorting mechanism that reflects the immune system’s plasticity.

Although the precise localization of IFNγ on SMAPs remains unclear, whether encapsulated within the particles or attached to their surface, deposits of IFNγ particles are consistently observed at the synaptic cleft following granule fusion (Figs. 3e,f, 4, and 5). Additionally, IFNγ puncta lacking defined SMAP markers constitute a significant population in IFNγ-mCherry expressing stimulated CD8⁺ T cells on supported lipid bilayers (Figs. 3e,f). These IFNγ⁺ vesicles might also reside on extracellular vesicles (EVs) as we observed from the distal membrane secretion (Fig.7). Given that IFNγ receptors have been identified on EVs, IFNγ can be carried by EVs to modulate immune responses^46^. Since IFNγ contributes to cytotoxicity, it may be reasonable to include IFNγ⁺ particles as SMAPs if these particles possess a glycoprotein shell structure similar to that of SMAPs.

Our results reveal distinct ’lytic’ and ’non-lytic’ IFNγ populations in activated effector T cells, comprising both pre-existing and newly synthesized IFNγ. While our data highlight the functional and spatial diversity of these IFNγ populations, it remains unclear whether, after initial synaptic depletion, the remaining ’lytic IFNγ’ repolarizes from the synapse to the distal membrane, or if newly generated ’lytic IFNγ’ directly polarizes to the distal membrane. Additionally, the exact cellular compartment responsible for packaging non-lytic IFNγ remains unclear. Our data suggest the possibility of multivesicular bodies (MVBs) as the source, given that IFNγ particles were released at the distal membrane site (Fig.7 and Extended Data Fig.5). Alternatively, they may originate from another source of IFNγ, such as small vesicles observed in TIRF imaging. Several studies have documented the widespread effects of IFNγ on antitumor immunity^30, 31, 38, 47^, suggesting a broader secretion pattern. This observation provides new insight to the canonical understanding that IFNγ are released exclusively at the synapse^29^. The absence of synaptic secretion in Munc13-4 KO cells and the observed distal membrane release highlight the adaptability of IFNγ secretion pathways, which may play a crucial role in navigating the complex tumor microenvironment and supporting prolonged immune responses.

In conclusion, we identified lytic IFNγ stored within cytotoxic granules and its co-release with GzmB at the immune synapse reveals a previously unrecognized mechanism of IFNγ storage and release. The potential distinct role of IFNγ in cytotoxicity and immune modulation, combined with its functional flexibility, offers new avenues for enhancing T cell-mediated tumor killing and may inform novel therapeutic strategies in cancer immunotherapy.

## Methods

### Mice

Wild-type (WT) mice were purchased from Charles River. Synaptobrevin2-mRFP knock-in (Syb2 KI)^35^, granzyme B-mTFP knock-in (GzmB KI)^33^, and Munc13-4 knockout (KO)^36^ mice were generated as described previously. All transgenic and WT mice used in this study were on a C57BL/6N background, except for GzmB KI mice, which were on a C57BL/6J background. Mice of both sexes, aged 15-22 weeks, were used for experiments. Animals were housed under standard conditions (22°C, 50–60% humidity, and a 12-hour light/dark cycle). All experimental procedures were conducted in compliance with regulations of the state of Saarland (Landesamt für Verbraucherschutz, AZ.: 2.4.1.1 and 11/2021).

### Human CTLs

Human peripheral blood mononuclear cells (hPBMC) were provided by the blood bank of University Hospital of the Saarland. The local ethics committee of the Saarland Medical Association (Ärztekammer des Saarlandes) approved the research with human material conducted for this study (Az. 98/15). Human CD8⁺ T cells were positively isolated from peripheral blood mononuclear cells (PBMCs) of healthy donors using the Dynabeads FlowComp Human CD8 Kit. CD8⁺ T cells were activated and cultured under the same conditions as described for mouse CTLs, with AIM-V medium supplemented with 10% FCS, 0.5% penicillin-streptomycin, and 100 U/mL human recombinant IL-2 (Gibco) to support expansion. Cells were cultured at 37°C in a 5% CO₂ atmosphere.

### Cell culture

Splenocytes were isolated from 15–22-week-old mice as described previously^21^. Naive CD8+ T cells were positively selected using the Dynabeads FlowComp Mouse CD8 Kit (Invitrogen), following the manufacturer’s protocol. Isolated CD8+ T cells were stimulated with anti-CD3/anti-CD28 activator beads (1:0.6 ratio) and cultured for 5 days in AIM-V medium (Invitrogen) supplemented with 10% FCS, 1% penicillin-streptomycin (Invitrogen), and 50 µM 2-mercaptoethanol. Cells were cultured at a density of ∼1 × 10⁶ cells/ml in 24-well plates. On day 2 post-stimulation, 100 U/ml recombinant IL-2 (Gibco) was added to support T cell proliferation. The resulting activated effector cytotoxic T lymphocytes (CTLs) were used for subsequent experiments.

For human CTL cultures, CD8+ T cells were positively isolated from peripheral blood mononuclear cells (PBMCs) of healthy donors using the Dynabeads FlowComp Human CD8 Kit (Invitrogen). CD8+ T cells were activated and cultured under the same conditions as described for mouse CTLs, with AIM-V medium supplemented with 10% FCS, 1% penicillin-streptomycin, and 100 U/ml human recombinant IL-2 (Gibco).

P815 target cells were cultured in RPMI medium (Invitrogen) supplemented with 10% FCS, 1% penicillin-streptomycin (Invitrogen), and 10 mM HEPES (Invitrogen). All cell cultures were maintained at 37°C with 5% CO₂.

### Plasmids and antibodies

To construct the pMax-mIFNγ-mCherry plasmid, the mouse IFNγ gene sequence was amplified from the cDNA of mouse CD8 ⁺ T cells via PCR. The forward primer 5’-GGTACCATGAACGCTACACACTGCATCTTG-3’ and the reverse primer 5’- GAATTCGCAGCGACTCCTTTTCCGCTTCCT-3’ were utilized, incorporating KpnI and EcoRI restriction sites, respectively. The pMax-GzmA-mCherry plasmid was employed as the vector and treated with KpnI and EcoRI to remove the granzyme A sequence. The PCR-amplified product was also treated with the same enzymes to generate complementary sticky ends. The resulting fragments were ligated to yield the final pMax-mIFNγ-mCherry construct. The pMax- hIFNγ-mNeonGreen plasmid was generated by amplifying the human interferon-gamma gene sequence from the cDNA of human peripheral blood mononuclear cells (PBMCs) by PCR. The amplification was performed using the forward primer 5’- ATGCCGGAATTCGCCGCCACCATGAAATATACAAGTTAT-3’ and the reverse primer 5’- ATGTATACGCGGATCCCTGGGATGCTCTTCGACC-3’, which contained EcoRI and BamHI restriction sites, respectively. The vector plasmid, pMax-mSrgn-mNeonGreen, was digested with EcoRI and BamHI to excise the serglycin sequence. The amplified IFNγ PCR product was digested with the same enzymes to generate compatible cohesive ends. The digested vector and insert were ligated using T4 DNA ligase, resulting in the construction of the pMax-hIFNγ- mNeonGreen plasmid. The sequences of both final vectors were validated by sequencing, which was performed using the respective forward and reverse primers by Microsynth Seqlab.

Antibodies used in this work are described in detail in Table 1.

### Flow cytometry

For flow cytometry analysis, 0.5 × 10⁶ day 4–5 effector CD8+ T cells were resuspended in D-PBS (Gibco) and incubated in the dark on ice for 30 minutes with the following cell surface markers: anti-CD44-PE, anti-CD62L-APC, and anti-mIFNγ-Alexa488 (Table 1). Day 5 human effector CD8+ T cells were treated with 5 µg/ml Brefeldin A (Sigma; B6542) for 2 hours before fixation and staining with anti-hIFNγ-Alexa594. Intracellular IFNγ staining was performed using the Transcription Factor Buffer Set (BD Pharmingen). Data acquisition was performed on a BD FACSAria III analyzer (BD Biosciences) using BD FACSDiva™ software. Flow cytometry data were analyzed with FlowJo v10.0.7 software.

### Killing assay

Day 3 effector WT or Munc13-4 KO CTLs (5 × 10⁴ cells) were co-cultured with p815 target cells at the indicated T cell–target ratios in 96-well U-bottom plates. P815 cells were pre-stained with 0.5 µM CellTrace Far Red (Invitrogen) for 10 minutes at RT and washed once with D-PBS before co-culture with T cells. Cells were cultured in RPMI-based medium for 3 hours in the presence of anti-mCD3 antibody (5 µg/ml) at 37°C with 5% CO₂. To evaluate the contribution of IFNγ in killing, cells were treated with anti-IFNγ (clone XMG1.2) to neutralize secreted IFNγ in the medium or recombinant mouse IFNγ (Abcam) to increase soluble IFNγ levels. Following incubation, cells were stained with 0.5 µg/ml propidium iodide (PI) and analyzed by flow cytometry. Dead target cells were identified as APC⁺ PI⁺ populations.

### Total internal reflection fluorescence (TIRF) microscopy

For secretion analysis, day 5 effector mouse CTLs (mCTLs) isolated from GzmB KI mice and day 5 human CTLs were used.

Mouse CTLs were transfected with mIFNγ-mCherry, while human CTLs were co-transfected with hIFNγ-mNeonGreen and hGzmB-mCherry via electroporation (Lonza kit). Cells were allowed to express transgenes for 12 hours at 32°C in OptiMEM-based transfection medium supplemented with 10% FCS, 10 mM HEPES, 1% DMSO, and 1 mM sodium pyruvate^48^ before TIRF microscopy. Two hours prior to imaging, activator beads were removed from the cultures. For imaging, 3 × 10⁵ cells were resuspended in 30 µl of extracellular buffer (10 mM glucose, 5 mM HEPES, 155 mM NaCl, 4.5 mM KCl, and 2 mM MgCl₂) and allowed to settle for 1–2 minutes on anti-CD3-coated coverslips (30 µg/ml; clone 2c-11 for mouse, UCHT1 for human). Cells were then perfused with extracellular buffer containing calcium (10 mM glucose, 5 mM HEPES, 140 mM NaCl, 4.5 mM KCl, 2 mM MgCl₂, and 10 mM CaCl₂) to stimulate cytotoxic granule (CG) secretion. The TIRF microscopy setup (Visitron Systems GmbH) was based on an IX83 microscope (Olympus) equipped with a UAPON100XOTIRF NA 1.49 objective (Olympus), solid-state excitation lasers at 488 nm, 561 nm, and 647 nm, an iLAS2 illumination control system (Roper Scientific SAS), an Evolve-EM 515 camera (Photometrics), and a filter cube containing Semrock FF444/520/590/Di01 dichroic and FF01-465/537/623 emission filters. The system was controlled using Visiview software (version 4.0.0.11, Visitron Systems GmbH). Images were acquired at a frequency of 10 Hz with 100 ms exposure time. Time-lapse series were analyzed using ImageJ or the FIJI package for ImageJ. CG secretion was quantified using ImageJ with the Time Series Analyzer plugin. A fusion event was defined as a fluorescence change within 300 ms accompanied by a diffusion cloud.

### Supported lipid bilayers (SLB)

SLBs were prepared as previously described ^49, 50^. Briefly, to prepare a clean glass chamber for SLB, the glass coverslips pre-washed with acid piranha and a plasma cleaner were mounted on sticky-Slide VI0.4 (Ibidi) to form 6 flow channels. Small unilamellar liposomes were prepared using 18:1 DGS-NTA(Ni) (790404C-AVL, Avanti Polar Lipids), 18:1 Biotinyl Cap (870282C-AVL, Avanti Polar Lipids), and 18:1 (D9-Cis) PC (850375C-AVL, Avanti Polar Lipids) in specific mixtures at total lipid concentration of 4 mM.

20 min at 20 ± 2°C. SLB were then washed with HEPES buffer containing 1 mM CaCl_2_ and 2 mM MgCl_2_ (HBS) and blocked with 1% human serum albumin (HBS/HSA) prior to functionalization. The SLB were functionalized in two steps. First with 5 µg per channel streptavidin (S11226, Thermo Fisher Scientific) for 10 min at 20 ± 2°C followed by three washings. Finally, 30 µg per channel biotinylated anti-mouse CD3ε (BD Pharmingen, clone 145-2C11) was linked to the streptavidin on SLB as well as 50 µg/ml 12-Histidine tagged mouse ICAM-1 was linked to Nickel ions.

### Confocal live cell imaging

Granule polarization to the synapse in CTLs targeting target cells was visualized using confocal microscopy (LSM 780, Zeiss) with 63x Plan-Apochromat objective (NA 1.4; Zeiss). Briefly, day 5 IFNγ-mCherry expressing GzmB KI CTLs (1 x 10^5^) were co-cultured with P815 target cells (2 x 10^4^) on poly-ornithine coated glass coverslip to ensure less cell movement for imaging. The RPMI co-culture medium contained 10 mM HEPES and 5 µg/ml anti-CD3ε antibody (Extended Data Table 1) to promote T cell to form contacts with target cells. Live imaging was performed under 37°C temperature control. Images were acquired as z-stacks with 1 Hz acquisition over time. The total thickness of the stack was 6 µm while distance between individual slices was 1 µm.

### Structured illumination microscopy (SIM)

For co-localization studies, T cells were fixed with ice-cold 4% paraformaldehyde (PFA) in D-PBS (Thermo Fisher Scientific) for 20 minutes, permeabilized with 0.1% Triton X-100 in D-PBS for 20 minutes, and blocked for 30 minutes in the same solution containing 2% bovine serum albumin (BSA). Cells were stained with anti- hIFNγ-Alexa594 (BioLegend, clone B27), anti-mIFNγ-Alexa647 (BioLegend, clone XMG1.2), and anti-GzmB-Alexa647 (BioLegend, clone GB11). For secreted SMAP detection on SLBs, 0.5 × 10⁶ T cells were seeded per ibidi channel and incubated for 90 minutes at 37°C with 5% CO₂ to allow synapse formation and SMAP secretion. Cells were washed five times with cold D-PBS to remove unbound cells while retaining deposited SMAPs. Deposited SMAPs were fixed with 2% pre-warmed PFA for 2 minutes at 37°C, washed three times with D-PBS, and stained with wheat germ agglutinin (WGA)-Alexa647 or WGA-Alexa488 (1 µg/ml) for 30 minutes at room temperature (RT). SMAPs were then permeabilized with 0.05% Triton X-100 for 5 minutes, blocked with 2% BSA for 30 minutes, and stained with anti-GzmB-Alexa647.

For analysis of IFNγ intracellular distribution and distal membrane secretion, WT and Munc13- 4 KO T cells transfected with IFNγ-mCherry, or human CTLs transfected with IFNγ-mNeogreen, were seeded on anti-CD3-coated glass coverslips for 30 or 60 minutes at 37°C with 5% CO₂. Cells were fixed with 4% PFA for 5 minutes and stained with WGA-Alexa488 (1 µg/ml) for 30 minutes at room temperature to mark the plasma membrane (PM). Deposited IFNγ on the coverslip was imaged to confirm distal membrane secretion. For quantification of the density of mouse IFNγ particles, cell-free areas were manually selected based on brightfield images using ImageJ. Particles were counted within the field of view containing transfected cells and normalized to the selected area (µm²). For human IFN-γ particle analysis, due to low transfection efficiency, the transfected cells were not in close proximity to each other in the images. Therefore, particles were manually counted within about 30 µm radius around each transfected cell. The analysis was reported as particles per cell. For IFNγ intracellular distribution, cells were washed twice with D-PBS, permeabilized, blocked, and stained with anti-GzmB-Alexa647. For T cell-target conjugates, 0.5 × 10⁴ T cells were mixed with calcein blue AM (Invitrogen)-labeled p815 target cells (1 × 10⁴) on polyornithine-coated coverslips in RPMI-based medium with 5 µg/ml anti-mCD3 (clone 2C11). Conjugates were fixed after 30 or 60 minutes with 4% PFA and stained with WGA-Alexa488 to mark the PM. For human CTLs, day 5 effector T cells were co-cultured with p815 target cells in the presence of 5 µg/ml anti- hCD3 (clone UCHT1), fixed after 1 or 2 hours, and stained with WGA-Alexa488. Following permeabilization and blocking, cells were stained with anti-hIFNγ-Alexa594 and anti-GzmB- Alexa647. Samples were mounted in Mowiol-based mounting medium. All experiments were performed on a Zeiss Elyra PS.1 microscopic SIM System with Zen 2010 software for device control and high-resolution image processing. Images were acquired as z-stacks with 0.2-µm intervals through the entire cell. Single-plane and maximum intensity projections were used to illustrate vesicle co-localization.

For correlative fluorescence and electron microscopy (CLEM) images of cells embedded in 100 nm ultrathin Lowicryl sections were acquired with excitation light of 405, 488 and 561 nm wavelengths. Almost the entire field of view of a 200-mesh grid (around 90 µm^2^) could be observed with a 63× Plan-Apochromat objective by SIM, allowing a perfect orientation relative to the grid bars. After adjusting the highest and lowest focus planes for z-stack analysis in brightfield, images were recorded with a step size of 100 nm. Fluorescent images were excited with 405 nm wavelengths to visualize DAPI. 488 nm wavelength was used for visualization of GzmB and 561 nm for IFNγ. The DAPI image (405 nm wavelength) was used to identify the nuclei, the plasma membranes of the CTLs and the image plane. 3-10 images were recorded with a step size of 100 nm to scan the cells of interest. All images were acquired with a 63× Plan-Apochromat (NA 1.4).

For fluorescence analysis of isolated cytotoxic organelles from Syb2-KI CTLs, centrifuged on gelatin coated coverslips from fraction 6 (MCGs) and 8 (SCGs), were permeabilized with 0.005% Triton-X-100 in Dulbecco’s PBS following blocking in PBS containing 2% BSA. Organelles were stained with anti-GzmB-Alexa647 and anti-IFNγ antibodies, while the secondary antibody was Alexa 488 (Extended Data Table 1). IFNγ, Syb2-mRFP and GzmB were visualized at 488 nm, 561 nm and 642 nm respectively.

### Cell homogenization and subcellular fractionation

Cytotoxic granule isolation was done as described^34^. 0.8-1.2 x 10^8^ activated CD8^+^ T cells from one Syb2 KI mouse, 0.7-0.8 x 10^8^ were harvested and washed once in buffer (Invitrogen, PBS with 0.1% BSA and 2 mM EDTA) before resuspension in 2 ml of homogenization buffer (300 mM sucrose, 10 mM HEPES (pH 7.3), 5 mM EDTA (pH 8.0) supplemented with protease inhibitors (3 mM Pefabloc, 10 µM E64 and 10 µg Pepstatin A). The cell suspension was transferred into a pre-chilled cell disruption bomb (Parr 1019HC T304 SS) connected to a nitrogen source. After 25 min equilibration at 800 psi the nitrogen pressure was released and the cell homogenate was collected. The cell lysate was centrifuged for 10 min at 1000 x g at 4°C to pellet unbroken cells, partially disrupted cells and nuclei. The resulting post-nuclear supernatant was layered on top of a discontinuous sucrose density gradient with 0.8, 1.0, 1.2, 1.4 and 1.6 M sucrose in 10 mM HEPES with 5 mM EDTA with protease inhibitors as described before (pH 7.3), 2 ml for each fraction. After ultracentrifugation at 100,000 x g for 90 min at 4°C in a SW40Ti rotor (Beckmann), twelve 1 ml fractions were collected from the top of the gradient and supplemented with fresh protease inhibitors. For fluorescence analysis, 1 ml each of fraction 6 and 8 was diluted in 10 ml D-PBS and centrifuged for 30 min at 10,000 x g at 4°C in a SW41Ti rotor (Beckmann) on gelatin-coated coverslips.

### RNA isolation, cDNA synthesis and RT-qPCR

WT or Munc13-4 knockout (KO) CD8⁺ T cells (1 × 10⁶ cells per well) were stimulated on anti-CD3-coated plates (30 µg/ml) or cultured on polyornithine (Sigma) -coated plates as an unstimulated control in a 24-well plate for 10 or 20 hours. Total RNA was extracted using TRIzol reagent (Thermo Fisher Scientific) following the manufacturer’s protocol. Briefly, 1 × 10⁶ CD8⁺ T cells were homogenized in 500 µl of TRIzol reagent, followed by the addition of 100 µl of chloroform. The samples were centrifuged at 12,000 × g for 15 minutes at 4°C, separating the mixture into three distinct phases, with the RNA remaining exclusively in the upper aqueous phase. The aqueous phase was carefully transferred to a fresh tube without disturbing the interphase, and the RNA was precipitated by mixing with an equal volume of 100% isopropanol. The RNA pellet was washed once with 75% ethanol, air-dried, and dissolved in DEPC-treated H₂O. The quality and quantity of the extracted RNA were assessed using a NanoDrop spectrophotometer.

The isolated RNA was subsequently reverse transcribed into complementary DNA (cDNA) using the ABScript II cDNA First-Strand Synthesis Kit (ABclonal) in accordance with the manufacturer’s instruction. Briefly, 1 µg of RNA was mixed with a reaction mixture containing Random Primer Mix, 10 mM dNTPs and ABScript II enzyme. The reaction was first incubated at 25°C for 5 minutes, followed by incubation at 42°C for 1 hour. Enzyme inactivation was achieved by heating the reaction to 80°C for 5 minutes. Real-time qPCR (RT-qPCR) was performed using Genious 2X SYBR Green Fast qPCR Mix (ABclonal) on a Bio-Rad CFX96 Touch Real-Time PCR Detection System. A total of 10 ng of cDNA was used as template in each reaction. Amplification was performed according to the manufacturer’s protocol using specific primers targeting Munc13-1 (forward: 5′-CAACTGGAATTACTTTGGC-3′, reverse: 5′- GCGAAGTCGTATAGTAGG-3′) and Munc13-4 (forward: 5′- GCCATTCTGCCCCTGATGAAAT-3′, reverse: 5′-ACCTCCACCAGCACAGTAAG-3′). TBP1 (Qiagen, QT00198443) was used as an internal reference gene to normalize target gene expression. Relative gene expression levels were quantified using the 2^(-ΔCt) method.

### ELISA

WT or Munc13-4 KO CD8⁺ T cells (1 × 10⁶ cells per well) were stimulated on anti-CD3- coated plates (30 µg/ml) in 24-well plates containing 1 ml AIM-V-based culture medium. Unstimulated control cells were plated on polyornithine-coated plates. Supernatants were collected at the indicated time points during 20 hours of incubation at 37°C. IFNγ concentrations were quantified using an IFNγ ELISA kit (Abcam, ab46081) according to the manufacturer’s instructions. Briefly, 100 µL of 1:10-diluted culture supernatants were added to IFNγ antibody-precoated 96-well plates and incubated for 1 hour to form antibody–antigen complexes. Following washes, 100 µL TMB substrate was added and incubated for 30 minutes at RT in the dark. The reaction was stopped with 50 µL of stop solution, and absorbance was measured at 450 nm using a TECAN Infinite M200 Pro plate reader. Four independent co-culture preparations were analyzed.

### Electron microscopy

Post-embedding correlative light and electron microscopy (CLEM) of cryo-fixed mouse CTLs was done as described previously^21^. Day 4 GzmB-mTFP KI CTLs were transfected with IFNγ-mCherry for 6-7 hours. Cells were seeded on 1.4 mm sapphire discs coated with poly-L-ornithine (0.1 mg/ml) and anti-CD3ɛ (30 µg/ml) in flat specimen carriers (Leica) and vitrified in a high-pressure freezing system (EM PACT2, Leica). The frozen specimens were cryo-transferred into the precooled (-130°C) freeze substitution chamber of the AFS2 (Leica). The temperature was increased from -130 to −90°C for 2h. Cryo-substitution was performed at −90°C to −70°C for 20 h in anhydrous acetone and at −70 to −60°C for 24 h with 0.3% (wt/vol) uranyl acetate in anhydrous acetone. At −60°C the samples were infiltrated with increasing concentrations (30, 60 and 100%; 1 h each) of Lowicryl (3:1 K11M/HM20 mixture; Electron Microscopy Sciences). After a 5 h 100% Lowicryl infiltration, samples were UV polymerized at −60°C for 24 h and for additional 15 h while temperature was raised linearly to 5°C. Samples were stored in the dark at 4°C until further processing. After removing the membrane carriers, 100 nm ultrathin sections were cut using an EM UC7 (Leica) and collected on carbon-coated 200 mesh copper grids (Plano). 1 d after sectioning a grid was stained with DAPI for 3 min (1/1000), washed and sealed between a coverslip and a glass slide for high resolution SIM imaging. After fluorescence imaging, the grid was carefully removed, stained with uranyl acetate and lead citrate and recorded with a Tecnai12 Biotwin electron microscope. Only CTLs with well conserved membranes, cell organelles and nuclei were analyzed and used for correlation. The DAPI signal of the nucleus was used to find the best overlap of the SIM image with the electron microscope image. The final alignment defines the position of the fluorescent signals within the cell of interest. Images were overlaid in Corel DRAW 2021.

### Imaging data and statistical analysis

All statistical tests were performed using Igor Pro or SigmaPlot 14.0, and data are presented as mean ± SEM (or SD where specified). The statistical tests used are indicated in the respective figure legends. Unless stated otherwise, two-group comparisons were performed using the Mann-Whitney U test. All pp-values were calculated with two-tailed statistical tests and a 95% confidence interval. Statistical significance was denoted as *p < 0.05, **p < 0.01, ***p < 0.001. All imaging data were analyzed with ImageJ or FIJI (version 15). Object-based co-localization analysis was conducted using the DiAna plugin in ImageJ (v52). Individual fluorescent spots were identified through an iterative segmentation procedure with the following parameters: step value of 100, size range of 15- 700 pixels, and independently adjusted thresholds for each channel.

## Data availability

Additional data and information that support the findings of this study are available from the corresponding author upon reasonable request.

## Acknowledgments

This work was supported by the China Scholarship Council (CSC), grants from the Deutsche Forschungsgemeinschaft (SFB 894 to J.R. and E.K.), European Commission (ERC -2021- SyG_951329 to J.R., S. V., C.T.B. and M.L.D) and University of Saarland (HOMFORexzellent2020 and NanoBioMed Young Investigator grant 2020 to H.-F.C.). We thank Margarete Klose, Anja Bergsträßer, Nicole Rothgerber and Katrin Sandmeier for excellent technical assistance.

## Author Contributions

V.P., E. K. and H.-F. C. conceived the study. X. L., M.-L. W., H.-F. C. and M. H. performed FACS experiments and analysis. M.-L. W., N. A. and H.-F. C. contributed to co-localization study. X. L., S.-M. T. and C.-H. L. performed ELISA and qPCR analysis. H.-F. C. and X. L. performed imaging experiments and analysis. C. S. performed CLEM experiments, organelle isolation, imaging and analysis. A.C. and U.B. developed the macros for object-based analysis.

J. R., E.K. and H.-F. C. acquired the funding. H.-F. C., E. K. and C. S. provided critical assistance in data interpretation. E. K. and H.-F. C. supervised the study. H.-F. C. wrote the manuscript; all authors edited and approved the paper.

## Additional information

**Extended Data Table 1.** Antibodies used in this study.

| Antibody<br>anti- | Supplier | Identifier | Source | Dilution | application |
| --- | --- | --- | --- | --- | --- |
| CD3ε | BD Pharmingen | 145-2C11 | Mouse | 1:33 | Glass<br>coating |
| Mouse IFNγ-Alexa488 | Biolegend | XMG1.2 | rat | 1:200 | FACS<br>ICC |
| Mouse IFNγ-Alexa647 | Biolegend | XMG1.2 | rat | 1:200 | ICC |
| Mouse IFNγ | Biolegend | XMG1.2 | rat | 1:50-1:400 | Neutralization<br>Killing assay |
| Granzyme B-Alexa647 | Biolegend | GB11 | Mouse | 1:100 | ICC |
| CD25-FITC | BD Pharmingen | 7D4 | Rat | 1:200 | FACS |
| CD44-PE | eBioscience | IM7 | Rat | 1:200 | FACS |
| CD62L-APC | BD Pharmingen | MEL-14 | Rat | 1:200 | FACS |
| Human IFNγ-Alexa594 | Biolegend | B27 | mouse | 1:200 | ICC<br>FACS |
| Human CD3 | Biolegend | UCHT1 | mouse | 1:33 | Glass<br>coating |

## Extended data

**Extended Data Fig. 1:**
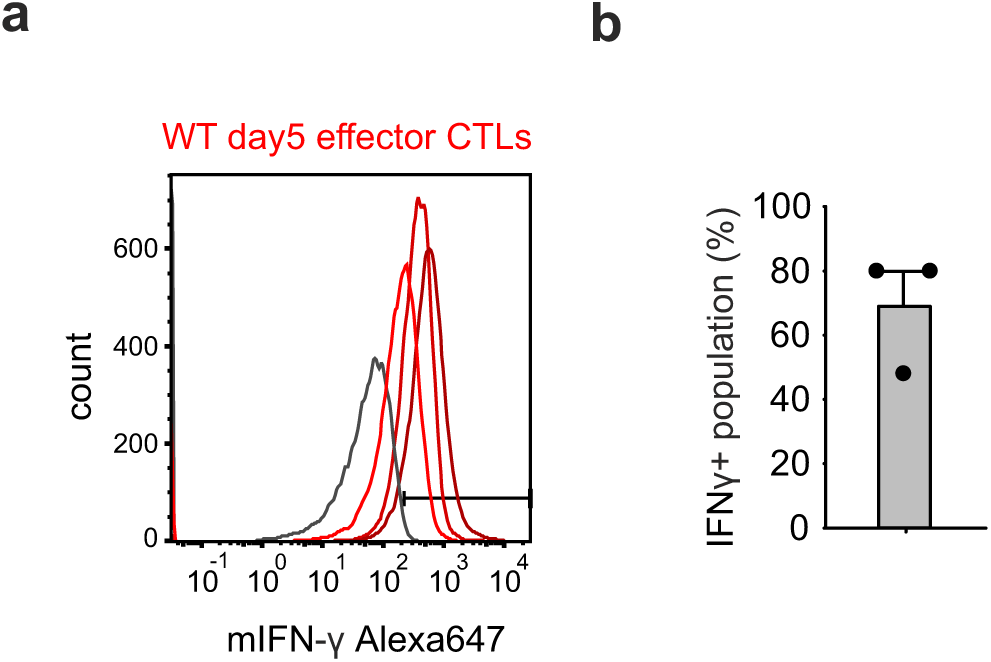
Day 5 mouse effector CTLs retain IFNγ expression without additional restimulation. (**a**) Flow cytometry analysis of WT day 5 effector cells without restimulation (N = 3). Histograms show unstained controls (gray line) and cells stained with anti-mIFNγ-Alexa647 from three preparations (red lines). (**b**) Quantification of IFNγ⁺ cell populations from (**a**).

**Extended Data Fig. 2:**
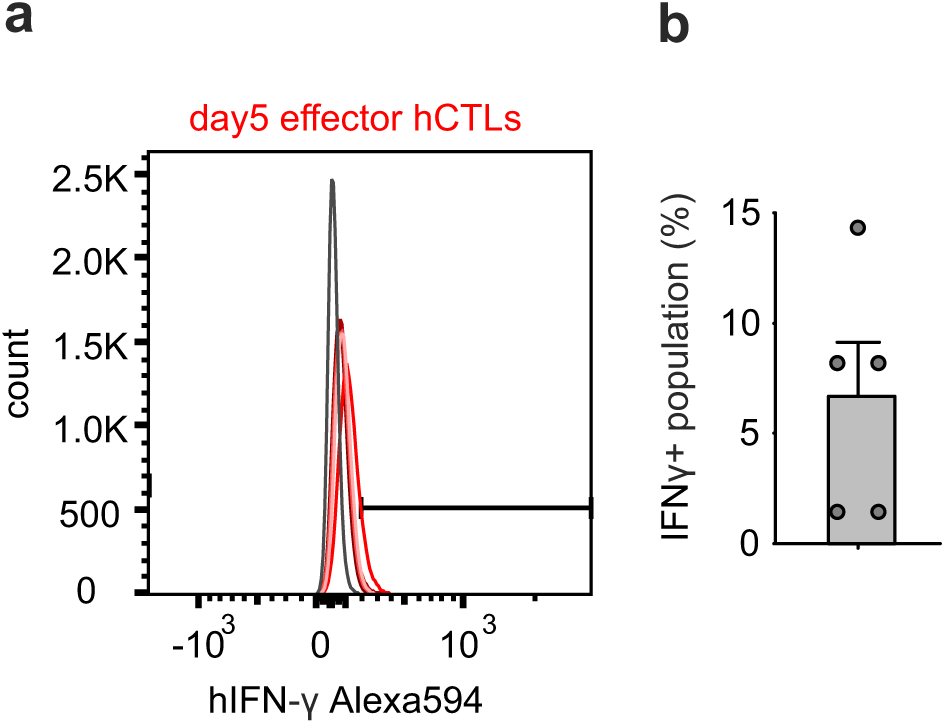
Day 5 human effector CTLs retain IFNγ expression without additional restimulation. (**a**) Flow cytometry analysis of day 5 effector CTLs from five donors. Cells were treated with Brefeldin A for 2 h, fixed, and stained with anti-hIFNγ-Alexa594. Histograms show unstained controls (gray line) and IFNγ-stained cells from 5 individual donors (red lines). (**b**) Quantification of IFNγ⁺ cell populations from (**a**).

**Extended Data Fig. 3:**
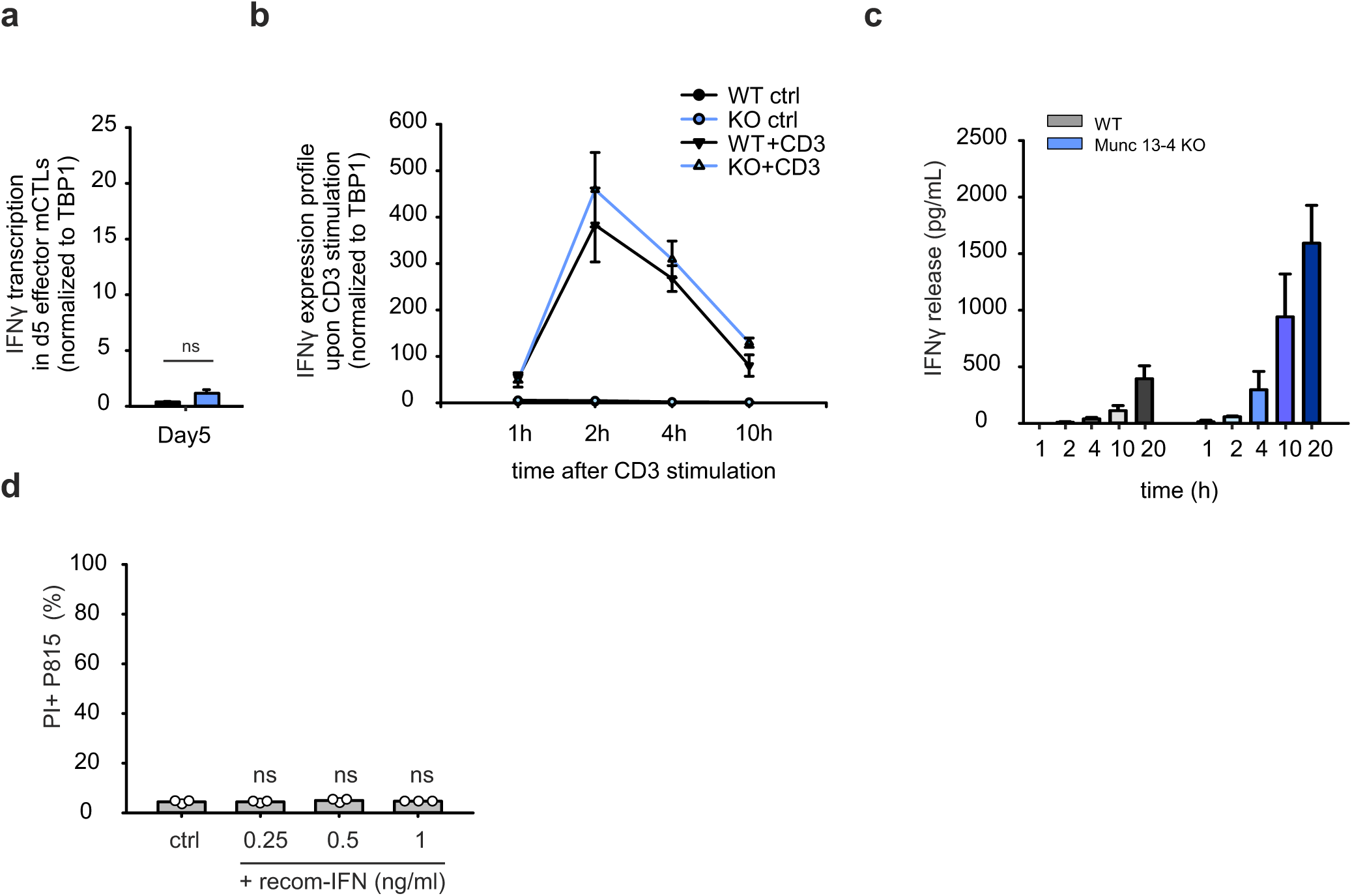
IFNγ mRNA expression, secretion, and functional analysis in WT and Munc13-4 KO CTLs. (**a**) Quantitative PCR (qPCR) was performed to analyze IFNγ transcription in WT and Munc13-4 KO day 5 effector CTLs (N=3). (**b**) IFNγ gene transcription in day 5 effector WT and Munc13-4 KO cells with or without CD3 restimulation. Cells were plated on polyornithine-coated (unstimulated control) or anti-CD3-coated plates for the indicated times within 10 hours. mRNA was extracted from these cells for qPCR analysis (N=3). (**c**) Constitutive release of IFNγ in WT and Munc13-4 KO effector CTLs. Day 5 effector cells were plated on polyornithine-coated plates. Supernatants from these cell groups were collected at the indicated time points within 20 hours. An ELISA assay was performed to measure the amount of released IFNγ (N=3). (**d**) P815 target cells were treated with mouse recombinant IFNγ to evaluate its cytotoxicity. Dead cells were stained with PI and analyzed by FACS (N = 3).

**Extended Data Fig. 4:**
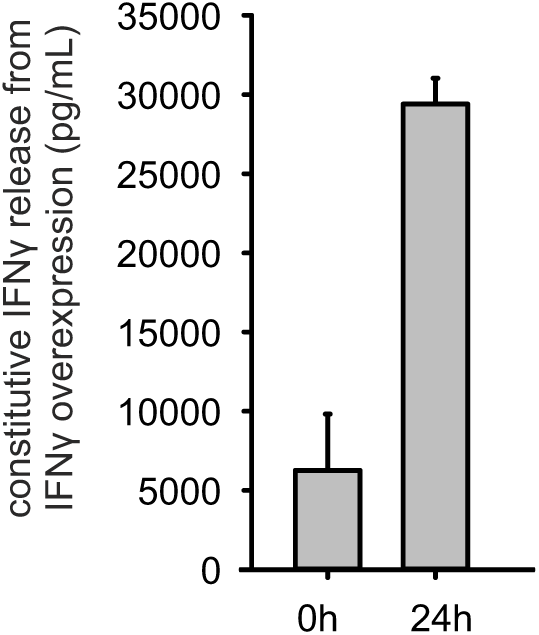
Overexpression of IFNγ in effector CTLs leads to increased constitutive secretion in the absence of CD3 stimulation. Day 5 effector WT cells were transfected with IFNγ-mCherry for 15 hours before the first supernatant collection (0 h). After 24 hours, the supernatant was collected again from the same culture. An ELISA assay was performed to measure the amount of released IFNγ. Data were collected from one independent experiment with three technical replicates.

**Extended Data Fig. 5:**
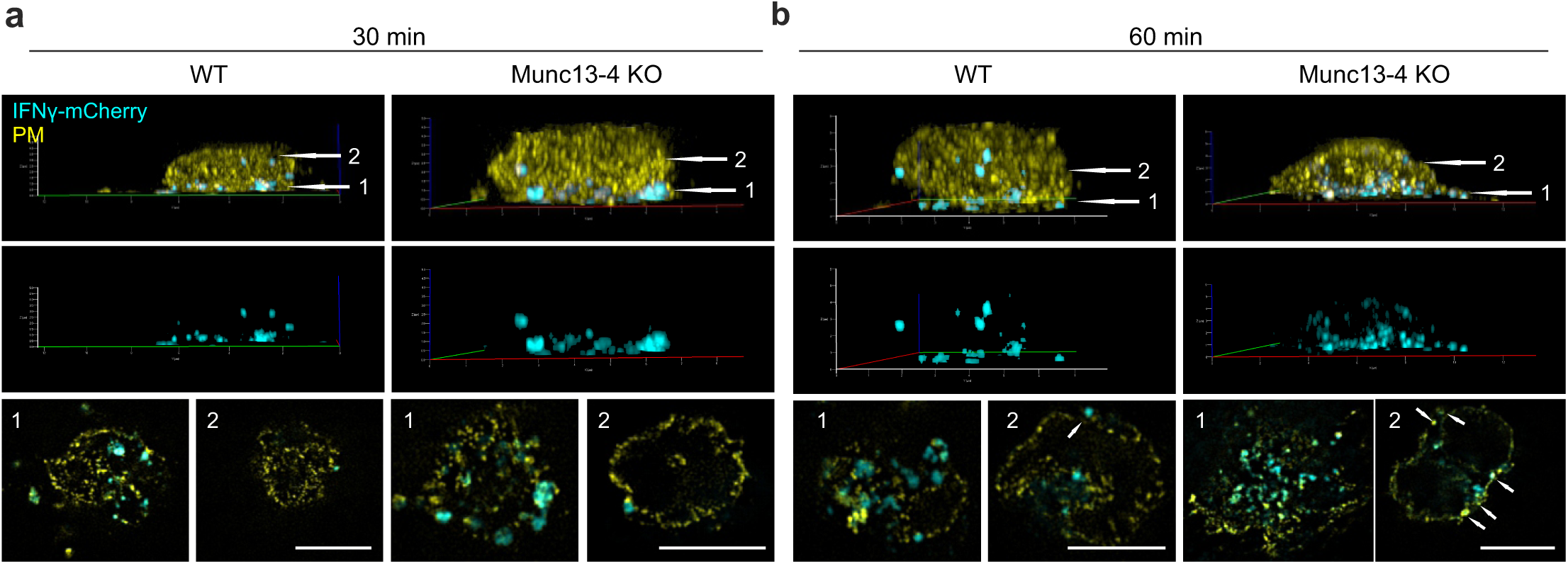
Polarization of IFNγ⁺ granules to the distal membrane in sustained synapses of mouse CTLs. (**a**, **b**) SIM images of day 5 effector WT and Munc13-4 KO CTLs fixed after 30 min (**a**) and 60 min (**b**) on anti-CD3-coated coverslips. CTLs were transfected with IFNγ-mCherry (cyan) and stained with WGA (yellow) to label the plasma membrane. 3D reconstructed images show lateral views of granule polarization at the synapse, with (upper panels) and without (middle panels) plasma membrane labeling. White arrows highlight IFNγ⁺ vesicles located at the synaptic layer (position 1) and distal membrane layer (position 2). Corresponding 2D xy images show single planes of the synaptic layer (position 1, lower left panel) and the distal membrane region (position 2, lower right panel). Small white arrows in the distal membrane region indicate IFNγ⁺ vesicles attached to the plasma membrane at 60 min. Scale bar, 5 μm.

